# Adaptive-optics lattice light-sheet microscopy enables correlative single-molecule imaging of endogenous proteins in native environments

**DOI:** 10.64898/2026.10.10.758183

**Authors:** Maxime Malivert, Chiara C. Galizia, Bram A.A.J.W. Willems, Magali Mondin, Cécile Lemoigne, Paul Lecompte Saint-Jean, Barthélémy Demeer, Maximilien Marcelino, Dario Crippa, Olivier Thoumine, Fabrice Harms, Matthieu Sainlos, Daniel Choquet, Mathieu Ducros, Angela M. Getz

## Abstract

Resolving molecular organization and dynamics in native environments remains challenging, as single-molecule imaging requires high sensitivity and spatiotemporal resolution while preserving 3D cellular context. Here, we present ALPINE (Adaptive-optics Lattice light-sheet microscopy for endogenous Protein Imaging in Native Environments), a modular framework for quantitative single-molecule imaging into complex biological samples. ALPINE combines adaptive-optics lattice light-sheet microscopy, programmable PSF shaping and acquisition stabilization, an engineered cell-specific labelling toolkit for endogenous cell-surface proteins and analysis pipelines assigning molecular localizations and trajectories to 3D cellular compartments. Labelling endogenous AMPA receptors in intact brain tissue, we demonstrate aberration-corrected 3D single-molecule localization and nanoscale organization within neuronal compartments. Importantly, fast optical sectioning of lattice light-sheet illumination further enables 3D single-molecule tracking in live tissue and quantification of molecular dynamics within native cellular environment. Together, ALPINE provides a modular approach to investigate how endogenous molecules are organized and dynamically regulated in complex biological systems, with components adaptable to diverse biological questions and imaging modalities.

## Introduction

Cellular functions emerge from molecular events dynamically organized within complex 3D environments, shaped by neighbouring cells, extracellular components and tissue architecture. Understanding these processes requires resolving how endogenous molecules are organized, interact and move within their native environment. Advances in genome editing and super-resolution microscopy increasingly enable proteins to be visualized at endogenous levels and beyond the diffraction limit. However, distinguishing individual molecules while maintaining compatibility with prolonged live imaging remains challenging, and single-molecule measurements of molecular organization and dynamics remain largely restricted to reduced-complexity experimental systems. Extending these measurements to native 3D environments is important for understanding how molecular behaviours integrate with cellular interactions and tissue architecture to give rise to emergent biological properties. Achieving this in living systems requires molecular sensitivity, high spatial and temporal resolution, compatibility with optically heterogeneous samples, and approaches that link molecular behaviour to its cellular context.

In neuroscience, this challenge is particularly relevant as nervous system function relies on specialized molecular organization and dynamics across diverse cell types and cellular compartments. Cell-surface proteins are central to many of these processes, mediating interactions between cells and their extracellular environment through receptors, adhesion molecules and signalling proteins. At synapses, differences in protein composition, nanoscale organization and mobility have emerged as key determinants of synaptic specificity, transmission and plasticity^1^. Synapse-specific regulation of AMPA receptor (AMPAR) biophysics and diffusion-trapping can tune synaptic integration and short-term plasticity^2^. Receptors at the post-synaptic density (PSD) are dynamically organized into nanoscale assemblies that regulate synaptic function^3,4^. Single-molecule localization microscopy (SMLM) and single-particle tracking (SPT) have revealed the nanoscale organization^5–7^ and mobility^8–10^ of individual receptors. However, these measurements have been performed predominantly in dissociated neuronal cultures, leaving how receptor nanoscale organization and dynamics vary across their native synaptic environments largely unexplored.

Lattice light-sheet microscopy (LLSM) provides an attractive foundation for single-molecule imaging in living 3D systems. With optimized optical sectioning, it combines high spatiotemporal resolution and detection efficiency with reduced photobleaching and phototoxicity, enabling prolonged live imaging from cultured cells to developing *C. elegans* and *Drosophila* embryos^11^. LLSM has further enabled single-molecule detection and tracking^11–13^, but these applications have largely focused on optically accessible samples. In heterogeneous tissues, such as the densely packed and lipid-rich brain, refractive-index variations increasingly distort the point spread function (PSF) with imaging depth, degrading resolution, signal to noise ratio (SNR)^14^ and single-molecule localization accuracy^15,16^. Nanoscale imaging in neuroscience has largely relied on approaches such as STED microscopy, which generally does not provide single-molecule localization and requires high photon exposure, or expansion microscopy, which requires fixed and physically expanded samples, leaving a need for approaches compatible with single-molecule measurements in living intact tissue. Combining LLSM with adaptive optics (AO) can compensate these aberrations and maintain imaging performance at depth, while programmable PSF engineering can encode the axial position of individual emitters for 3D localization^17–19^.

Beyond optical challenges, single-molecule imaging of endogenous proteins requires labelling strategies that combine molecular specificity with minimal perturbation. Probe size, linkage error, labelling density and fluorophore properties directly affect molecular measurements^20^. Small genetically encoded tags combined with high-affinity probes can minimize linkage error while providing access to bright, photostable fluorophores compatible with single-molecule imaging^21,22^. Cell-surface proteins are particularly amenable to this approach, as extracellularly accessible tags can be selectively targeted without intracellular delivery or membrane permeabilization, facilitating labelling in living samples while preserving cellular integrity^22–24^. In tissue, cell-specific labelling further enables endogenous proteins to be resolved within defined cells despite their expression in the surrounding tissue. Conditional strategies, such as cell-specific biotinylation of extracellular AP-tagged proteins combined with engineered avidin probes, therefore provide a versatile route for imaging endogenous cell-surface proteins in complex biological environments^25,26^. Beyond detecting individual molecules, understanding their behaviour requires considering how their organization and mobility are regulated by the local cellular environment. At synapses, interactions with scaffold proteins enriched at the PSD contribute to receptor anchoring and stabilization, regulating their nanoscale organization and mobility. Relating these molecular measurements to their cellular context is therefore essential for understanding how receptor behaviour varies across specialized neuronal compartments^4,27^. Correlative morphological imaging and segmentation can assign individual localizations and trajectories to defined cellular compartments, linking molecular measurements to cellular 3D architecture^28,29^.

Here, we present ALPINE (Adaptive-optics light-sheet Lattice microscopy of endogenous Proteins In Native Environments), an integrated and modular platform for quantitative imaging of endogenous protein organization and dynamics at the single-molecule level in complex biological environments (Supplementary Figure 1). We demonstrate ALPINE by imaging endogenous cell-surface proteins in organized brain tissue, combining (1) an AO-LLSM framework integrating automated aberration correction, programmable PSF shaping and acquisition stabilization for 3D single-molecule imaging at depth, (2) an engineered enhanced monovalent StreptAvidin (eSA) based labelling toolkit for cell-specific visualization of endogenous cell-surface proteins, optimized for single-molecule imaging in tissue and (3) analysis pipelines that assign single-molecule localizations and trajectories to segmented 3D cellular compartments, enabling compartment-resolved measurements of molecular organization and dynamics. Applying this framework to AP-tagged GluA2-AMPAR subunit and Neuroligin-1 (NLGNN1) reveals distinct molecular behaviours across cell-surface proteins, demonstrating its applicability to molecular targets with different properties. These modules can be combined into a complete workflow or implemented independently according to the biological question, providing adaptable strategies across molecular targets, cell types and biological systems. Together, ALPINE provides a versatile framework for investigating endogenous molecular organization and dynamics within native 3D environments.

## Results

### Adaptive-optics lattice light-sheet microscopy enables 3D single-molecule imaging in tissue

Single-molecule imaging in intact tissue, particularly under live-imaging conditions, requires an optical system combining efficient fluorescence detection and optical sectioning with high spatiotemporal resolution, while maintaining imaging performance in optically heterogeneous samples. To address these requirements, the ALPINE imaging framework builds upon a custom LLSM originally developed by Janelia Reasearch Campus^11^ and previously modified in our laboratory to include AO in the detection path for aberration correction^14^. To maintain diffraction-limited imaging at depth in intact brain tissue, we developed an Active Image Optimization (AIO) workflow that compensates for sample-induced optical aberrations. AIO consists of two sequential image-based optimization steps (Fig. 1a). First, an automatic light-sheet autofocus restores the overlap between the excitation plane and the detection focal plane, compensating for excitation defocus induced by refractive-index mismatch^30^. Second, a sensorless AO optimization iteratively applies low-order Zernike modes to a deformable mirror (DM) and maximizes an image-quality metric to recover the optimal wavefront. This strategy restores lateral and axial resolution without requiring guide stars or wavefront sensing during imaging, enabling near diffraction-limited imaging of fine neuronal structures at depth within brain tissue.

**Fig. 1.**
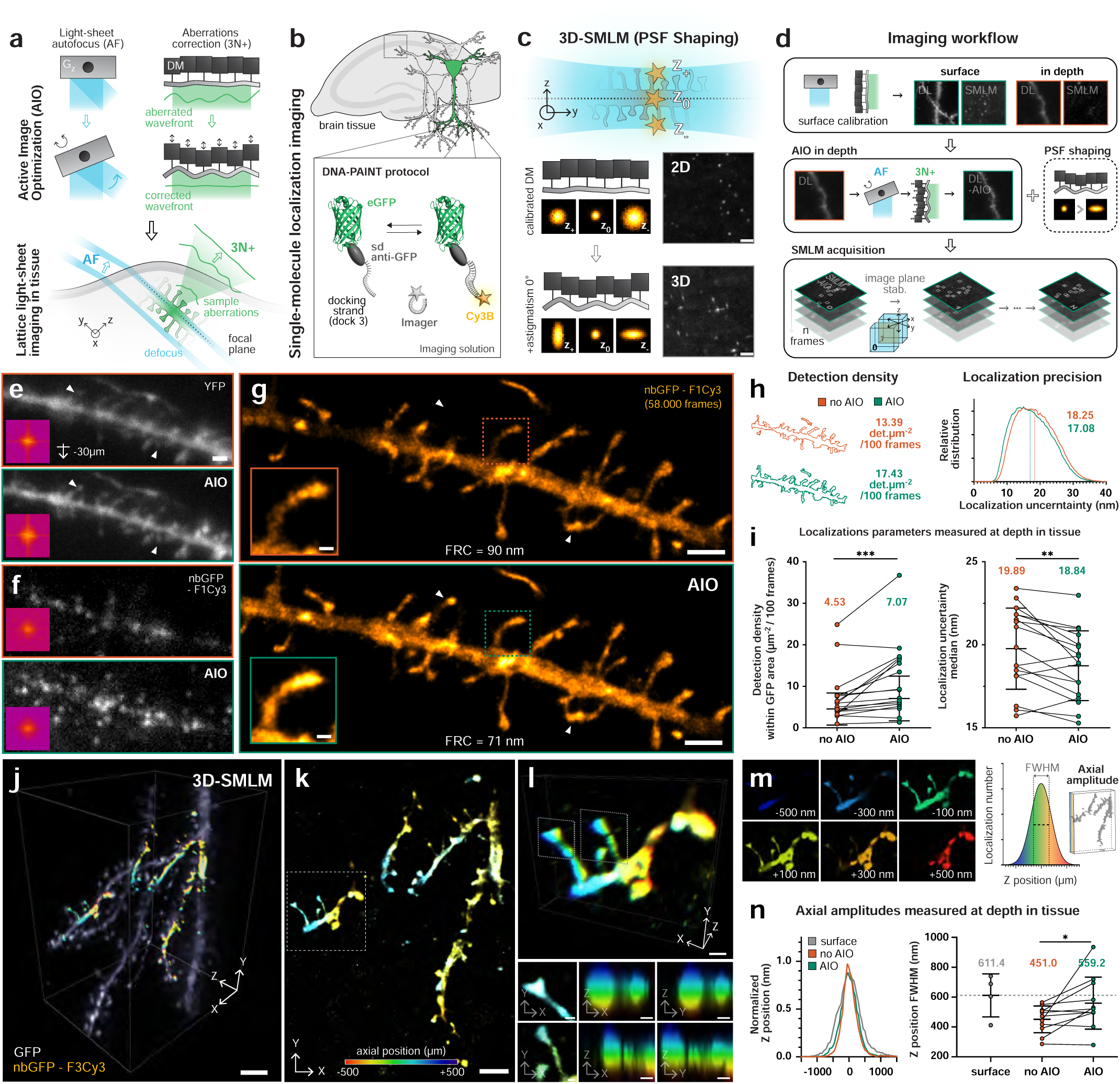
AO-LLSM for 3D-SR reconstruction imaging within tissue. **a,** Illustration of the AO-LLSM imaging in tissue with AIO correction procedure, comprising light-sheet AF and optical aberrations correction using the axial galvanometer (Gz) and deformable mirror (DM), respectively. **b,** Schematic of SMLM labelling strategy for brain tissue. Sparse and specific labelling of cells with eGFP enables DNA-PAINT imaging with low background and tuneable imaging wavelengths. **c,** Principle of 3D-SMLM through PSF shaping by applying controlled astigmatism 0° using the DM. **d,** Imaging workflow for SMLM in tissue. Top, Gz and DM calibrations are performed at the sample surface. Diffraction-limited (DL) and SMLM images are resolved at surface but aberrated in depth. Middle, AIO is applied to an in-depth field of view (FOV) to correct light-sheet defocus and optical aberrations on the DL channel, followed by PSF shaping for 3D-SMLM. Bottom, SMLM acquisition is performed as a series of sub-acquisitions interleaved with image-plane stabilization for drift correction. **e,** Representative MIPs of eGFP stacks acquired 30 μm below the tissue surface before (top) and after (bottom) AIO correction, with the corresponding power spectra. **f,** Average-intensity images (10 frames) from DNA-PAINT acquisitions using nbGFP–F1Cy3 imager in the same FOV as in **e**, before (top) and after (bottom) AIO correction, with the corresponding power spectra. **g,** SMLM reconstructions of the acquisitions shown in **f**, with the corresponding Fourier ring correlation (FRC) resolutions. Insets and arrows highlight improvements in the resolution and intensity of dendritic spines following AIO correction. **h,** Localization parameters from the acquisitions shown in **g**, illustrating the increase in the number of detected localizations within the eGFP-defined morphological signal and the reduction in median localization uncertainty following AIO correction. **i.** Comparison of localization density (normalized at 100 frames) within the eGFP-defined area and median localization uncertainty before and after AIO correction across acquisitions performed 20–50 μm below the tissue surface (700 frames per acquisition, N = 17, **P ≤ 0.005, ***P ≤ 0.0005, Wilcoxon test, paired t-test). **j,** 3D rendering of the eGFP morphological signal (grey) and 3D-SMLM reconstruction (color-coded by z position) within a single imaging plane. **k,** MIP of the 3D-SMLM reconstruction shown in **j**. **l.** 3D rendering of inset in **k**, with (bottom) associated MIPs XY, XZ and YZ of highlited spines. **m,** Combination of light-sheet thickness and PSF shaping enables localization detection over an axial range of approximately 1 μm. **n,** Left, normalized distributions of localization z positions measured at the tissue surface and at depths of 20– 50 μm. Right, corresponding full-width at half-maximum (FWHM) values for individual acquisitions (700 frames per acquisition, N_surface_ = 4, N_noAIO/AIO_ = 11, *P ≤ 0.05, paired t-test). **Scale bar,** 3 μm (**j**), 2 μm (**k, e, f**), 1 μm (**c, g**), 500 nm (**l-top**), 200 nm (**g-inset, l-bottom**).

To establish single-molecule imaging with AIO-LLSM in tissue, we used brain slices from Thy1-YFP knock-in mice, in which sparse neuronal YFP expression provides a morphological reference for imaging and subsequent single-molecule analysis. Because GFP-targeting nanobodies also recognize YFP, we used DNA-docking-strand-conjugated GFP nanobodies to implement DNA Points Accumulation for Imaging in Nanoscale Topography (DNA-PAINT) imaging of the labelled neuronal morphology (Fig. 1b)^31^. DNA-PAINT was particularly suited to our imaging configuration because transient imager binding provides sustained single-molecule blinking with high photon yields, while avoiding the oxygen-scavenging buffers typically required for Stochastic Optical Reconstruction Microscopy (STORM)-based approaches and their practical constraints in the LLSM sample chamber. In addition, the same GFP nanobody can be conjugated to different docking strands, including conventional and speed-optimized DNA-PAINT sequences, allowing binding kinetics, acquisition speed and localization performance to be adapted to experimental requirements^32,33^.

To extend single-molecule localization to 3D, we next exploited the DM for programmable PSF shaping^17^. Following AIO correction, a calibrated 0.1 µm root mean square (RMS) astigmatism was superimposed onto the corrected wavefront to encode the axial position of individual emitters within their PSF (Fig. 1c), enabling 3D single-molecule localization while maintaining aberration correction, which enables better axial localization precision as compared to the sole use of a cylindrical lens.

We integrated these components into a single acquisition workflow for 3D single-molecule imaging in tissue (Fig. 1d). AIO is first performed on the diffraction-limited morphological channel at the selected field of view (FOV), within a local isoplanatic patch^34^, defined to match the region where the light sheet maintains nearly constant thickness and spatial resolution^35^, ensuring homogeneous illumination and aberration correction across the imaged volume. PSF shaping is subsequently applied for 3D localization before switching to DNA-PAINT acquisition. Single-molecule imaging is performed as consecutive acquisitions interleaved with morphological imaging, enabling periodic axial drift correction through image-plane stabilization (Fig. S2).

Before evaluating AIO on native tissue aberrations, we first validated its performance under controlled conditions by introducing known aberrations with the DM (Fig. S3). Random combinations of low-order Zernike modes were applied during surface imaging, and successive AIO iterations corrected the induced wavefront aberrations and restored single-molecule localization performance. After three optimization cycles, the residual wavefront RMS decreased below the Maréchal criterion for diffraction-limited imaging at 560 nm (∼40 nm RMS), while localization uncertainty and detection yield returned to values comparable to the unaberrated condition (Fig. S3).

We next evaluated AIO under native tissue aberrations using diffraction-limited and DNA-PAINT acquisitions performed between 20 and 50 µm imaging depth in organotypic hippocampal slices from Thy1-YFP mice. In diffraction-limited YFP images, AIO produced modest changes in overall fluorescence intensity but improved the visibility of fine dendritic structures (Fig. 1e, white arrowheads), with Fourier transforms revealing a recovery of high spatial frequencies after correction. The effect was more pronounced for DNA-PAINT acquisitions (Fig. 1f), where AIO restored the single-molecule PSFs, increasing peak intensity and signal-to-background ratio. Consequently, SMLM reconstructions exhibited improved structural continuity and finer dendritic details, with Fourier ring correlation (FRC) resolution improving from 90 nm to 71 nm (Fig. 1g).

We next quantified these improvements using the localization statistics generated during SMLM reconstruction. Following AIO, the detection density, measured within the YFP-defined morphology, increased from 13.39 to 17.43 detections µm⁻² per 100 frames, while the median localization uncertainty decreased from 18.25 nm to 17.08 nm (Fig. 1h). These improvements are consistent with the restoration of a near-diffraction-limited PSF, concentrating emitted photons within the PSF core and thereby increasing single-molecule detection efficiency and localization precision.

To assess the robustness of these improvements, localization statistics were quantified across 17 independent acquisitions performed between 20 and 50 µm imaging depth (Fig. 1i). Consistent with the representative dataset, AIO significantly reduced the median localization uncertainty from 19.89 nm to 18.84 nm (paired t-test, P < 0.01), while increasing the detection density from 4.53 to 7.07 detections µm⁻² per 100 frames (paired t-test, P < 0.001). Together, these results demonstrate that AIO improves single-molecule imaging performance at depth, providing the optical conditions required for volumetric single-molecule imaging within brain tissue.

We next applied this engineered PSF to perform 3D DNA-PAINT imaging in tissue. This enabled 3D reconstruction of neuronal morphology within the light-sheet imaging volume, visualized as volumetric renderings and z-colour-coded projections (Fig. 1j–l). The astigmatism amplitude was selected to encode emitter positions across the ∼1 µm effective axial imaging range defined by the combined light-sheet excitation and detection volumes. Although the excitation profile has an axial full width at half maximum (FWHM) of ∼500–700 nm, the detection depth of field (∼0.7 µm)^14^, enables single-molecule events within the axial tails of the light sheet to remain detectable, extending the accessible localization range along z.

To experimentally quantify this accessible axial localization range, we measured the distribution of localization positions along z and determined its FWHM as the axial localization range (Fig. 1m). We then assessed whether this range was maintained at depth in tissue. Between 20 and 50 µm imaging depth, sample-induced aberrations compressed the axial distribution, reducing its FWHM from 611.4 nm at the tissue surface to 451.0 nm without AIO (Fig. 1n). This compression likely reflects reduced detection of emitters toward the axial extremes, where astigmatic PSFs become increasingly elongated and the associated photon loss is exacerbated by tissue-induced aberrations. AIO significantly increased the axial localization range to 559.2 nm, approaching the value measured at the tissue surface. These results demonstrate that AIO mitigates the compression of the axial localization range induced by tissue aberrations, preserving the astigmatic encoding required for 3D single-molecule localization at depth.

#### Biotin-Avidin labelling toolkit development for endogenous receptor imaging in tissue

Beyond the optical framework, the second component of ALPINE addresses the labelling requirements for single-molecule imaging of endogenous proteins in tissue: sparse and specific molecular labelling while preserving a morphological signal to identify the cellular context of individual molecules. To address these requirements, we focused on GluA2-containing AMPARs and employed the recently developed AP-GluA2 knock-in mouse line^25^, enabling cell-specific visualization of endogenous surface receptors within GFP-labelled neuronal morphology (Fig. 2a). In this approach, a short extracellular biotin acceptor peptide (AP) was introduced into the endogenous GluA2 subunit by CRISPR–Cas9-mediated genome editing. Cell-specific expression of the ER-retained biotin ligase BirAER catalyzes biotinylation of the AP tag, allowing surface receptors to be selectively labelled with fluorescent monovalent streptavidin (mSA)^24^, while co-expressed GFP reports BirA^ER^ expression and provides a cell volume marker for the corresponding neuronal morphology. More broadly, CRISPR-based endogenous tagging strategies in neurons provide a route to introduce extracellular AP tags into cell-surface proteins, extending this cell-specific biotinylation and streptavidin-based labelling strategy beyond GluA2 and NLGN1^36,37^. Labelling density can be tuned through the concentration of fluorescent mSA, enabling the sparse conditions required for single-molecule imaging while preserving endogenous receptor expression. However, the original mSA probe exhibited limited fluorescence intensity together with elevated non-specific background, restricting its application to single-molecule imaging in tissue and motivating the development of optimized biotin-binding probes.

**Fig. 2.**
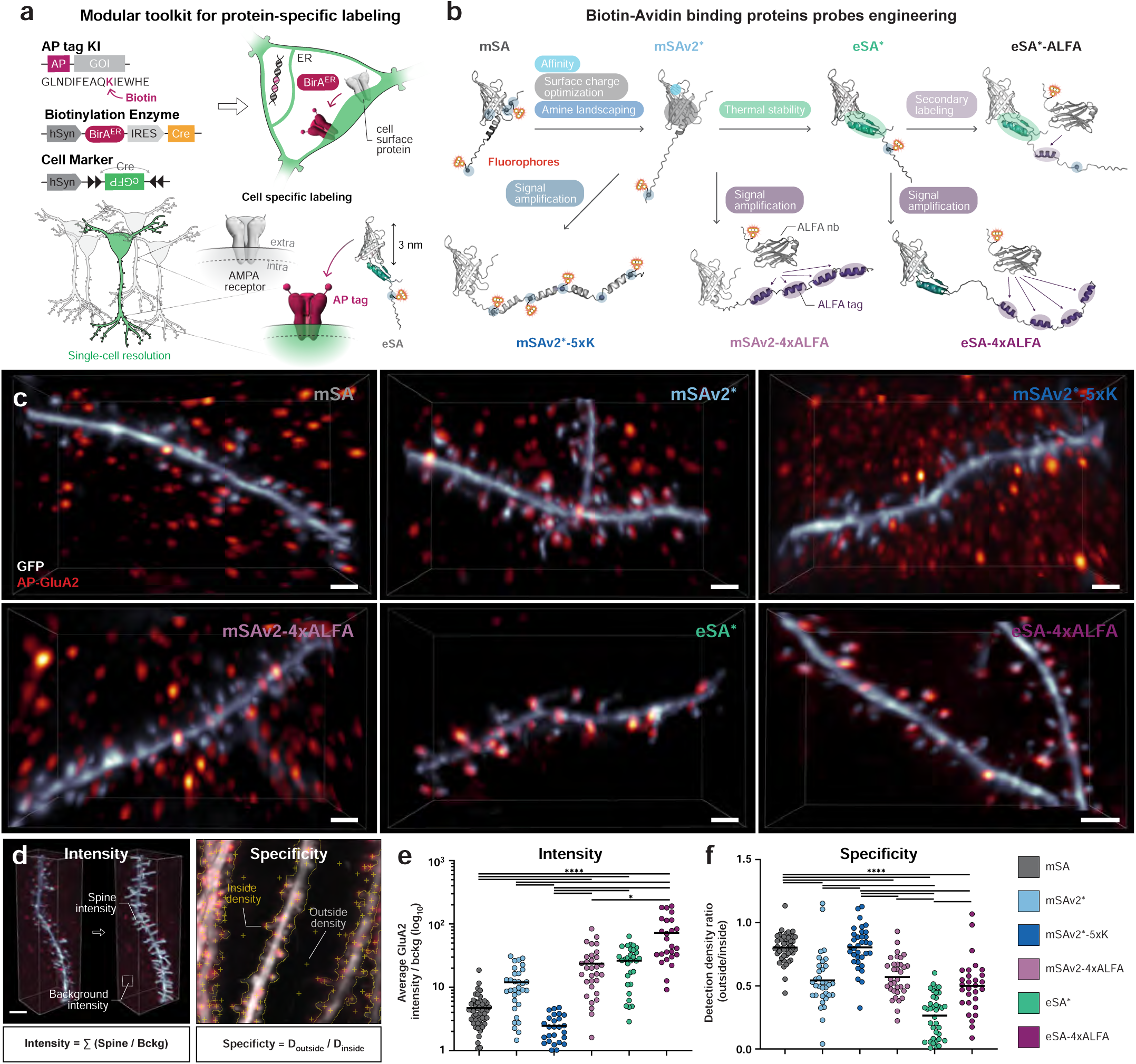
Avidin probes engineering for sparse and cell-specific visualization of endogenous proteins in tissue. **a,** Schematic representation of functionalized AP-GluA2 KI toolkit strategy for tuneable and cell-specific labelling of endogenous AMPARs. Targeted expression of ER-retained biotin ligase (BirAER) induces biotinylation of the extracellular AP tag allowing surface receptors to be visualized with small, high-affinity biotin-avidin (eSA) probes, linked to a fluorophore, at single-cell and synaptic resolution. **b,** Schematic representation of the monomeric streptavidin probes engineering strategy for tissue imaging. Biotin-binding proteins were sequentially engineered to improve affinity, surface charge, thermal stability and signal amplification, while introducing secondary labeling strategies for flexible and high-SNR visualization of endogenous proteins in fixed and lived tissue. Amino groups available for NHS-ester dye conjugation are depicted as blue spheres, with their number indicated in brackets. An asterisk stands for the presence of a single amino group. **c,** Representative LLSM imaging of dendritic spines in AP-GluA2 KI tissue with corresponding biotin-avindin probes defined in **b. d,** Schematic of custom analysis to measure the intensity and the specificity of each biotin-avidin probe. **e,** Comparison of the average AP-GluA2 signal normalized on the background (log10) for the biotin-avidin illustrated probes in **b** (N_mSA_ = 48, N_mSAv2*_ = 32, N_mSAv2*-5xK_ = 26, N_mSAv2-4xALFA_ = 31, N_eSA*_ = 30, N_eSA-4xALFA_ = 25, *P ≤ 0.05, ****P ≤ 0.0001, Kruskal-Wallis test). **f,** Comparison of the detection density ratio for the biotin-avidin illustrated probes in **b** (N_mSA_ = 44, N_mSAv2*_ = 34, N_mSAv2*-5xK_ = 33, N_mSAv2-4xALFA_ = 34, N_eSA*_ = 37, N_eSA-4xALFA_ = 29, ****P ≤ 0.0001, one-way ANOVA test). **Scale bar,** 2μm (**c**).

To systematically address the properties limiting endogenous receptor imaging in brain tissue, we engineered a series of monovalent biotin-binding probes with enhanced performance (eSA) (Fig. 2b). These limitations are particularly critical for single-molecule imaging, where probe brightness directly affects molecular detection and localization precision, while non-specific labeling can generate background events indistinguishable from individual target molecules. Starting from the original mSA, successive probe designs targeted optimization of biotin affinity, dye coupling control and regiospecificity, fluorescence intensity and labeling modularity. The first generation, mSAv2, incorporated optimization of the biotin-binding interface, amino group redistribution and surface charge to improve affinity while ensuring basic solubility and homogeneity following dye coupling via NHS-ester. Signal amplification was then explored through either increased fluorophore loading using a poly-lysine linker (mSAv2-5×K) or the introduction of four ALFA tag^38^ epitopes for secondary nanobody labeling (mSAv2-4×ALFA). Finally, thermostability improvement by combining computational optimization and fusion of both N- and C-termini to thermally stable domain yielded the optimized eSA, which was subsequently combined with the 4×ALFA tag module to generate eSA-4×ALFA. Beyond fluorescence signal amplification, the 4×ALFA tag module provides a versatile interface for secondary labeling, enabling direct compatibility with commercial ALFA tag-based DNA-PAINT reagents as well as straightforward adaptation to other ALFA tag-compatible labeling strategies.

We next evaluated whether these engineering steps translated into improved labelling performance and specificity in organotypic hippocampal slices using dye-conjugated probes and quantitative 3D LLSM imaging. Optimization of the streptavidin scaffold produced a marked increase in receptor fluorescence intensity together with a substantial reduction in non-specific labelling, both of which were readily apparent in the 3D LLSM images (Fig. 2c). To quantify these improvements, we measured fluorescence relative to background and labelling specificity from the ratio of detection densities outside and inside the GFP-defined neuronal morphology (Fig. 2d). Quantitative comparison across probes confirmed improvements in both parameters (Fig. 2e, f). Compared with the original mSA, optimization of the biotin-binding interface in mSAv2 increased the median receptor fluorescence by ∼2.8-fold, while subsequent thermostability engineering further enhanced the signal to ∼6.3-fold with eSA. Secondary labelling of eSA-4×ALFA with fluorescent ALFA nanobodies produced the highest fluorescence, reaching an ∼17-fold increase over mSA, while maintaining a consistent labelling pattern within the neuronal morphology. In parallel, labelling specificity improved with optimization of the streptavidin scaffold, as reflected by a reduction in the outside-to-inside GFP-defined morphology of BirAER-expressing neurons, in which AP-GluA2 is selectively biotinylated (Fig. 2f). eSA exhibited the lowest ratio, indicating the highest labelling specificity, whereas eSA-4×ALFA maintained similarly low background despite the additional secondary labelling step. Together, these results identify two complementary probe labelling strategies for distinct single-molecule applications. eSA combines high fluorescence intensity and labelling specificity with a minimal probe architecture and was therefore selected for SPT, whereas eSA-4×ALFA provides a modular secondary-labelling interface for signal amplification and DNA-PAINT while retaining the high specificity of the engineered scaffold.

#### ALPINE for 3D organization imaging of endogenous proteins in tissue

A central goal of ALPINE is to resolve and quantify the nanoscale organization of endogenous proteins within their native cellular environment. Because molecular assemblies are organized within three-dimensional cellular structures, incorporating axial information can provide complementary spatial information on their nanoscale organization and relationship to the surrounding cellular architecture organization^22^. Building on the optical and labelling developments described above, we next extended SMLM measurements to 3D to capture this additional dimension of protein nano-organization in tissue. AMPARs provide a particularly relevant model for this purpose, as they are organized into nanoscale clusters within the PSD, which can contain multiple higher density nanodomains. Super-resolution imaging in cultured neurons has shown that PSD typically spans several hundred nanometres laterally, while individual AMPAR nanodomains measure approximately 70–100 nm in diameter^6,7^.

To quantitatively resolve this 3D organization, we integrated endogenous GluA2 labelling, 3D DNA-PAINT imaging and point-cloud analysis into a unified workflow (Fig. 3a). The optimized eSA labelling toolkit enabled sparse and specific labelling of endogenous GluA2 in brain tissue, while programmable PSF shaping encoded the axial position of individual emitters during DNA-PAINT acquisition. Localization coordinates were reconstructed as 2D or 3D point clouds according to the acquisition mode, enabling direct comparison of receptor nano-organization with and without axial information.

**Fig. 3.**
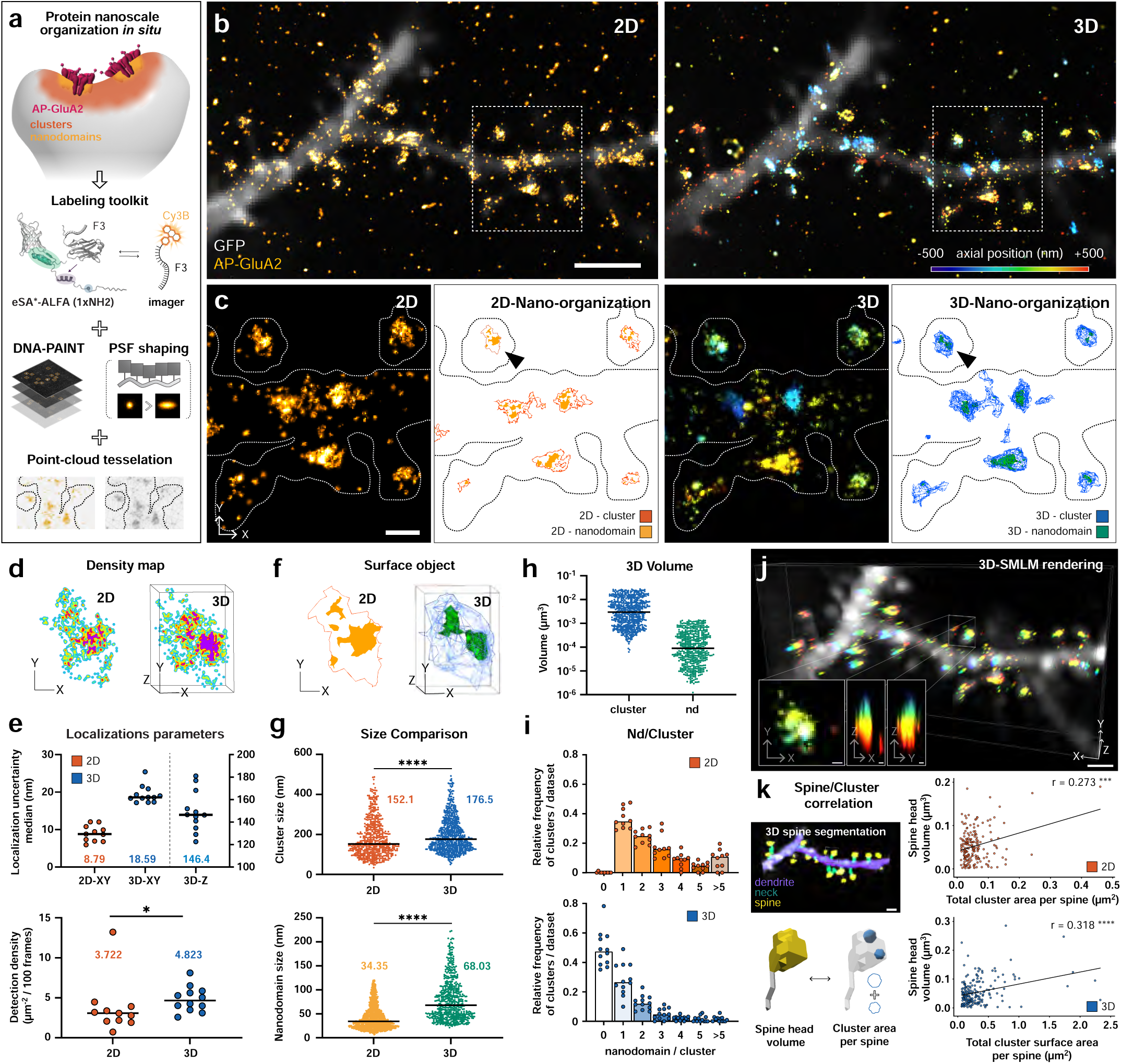
SMLM-AO-LLSM and biotin-avidin labelling toolkit to decipher *in situ* protein nanoscale organization in fixed brain tissue. **a,** Schematic representation of the imaging pipeline using biotin-avidin strategy adapted for DNA-PAINT imaging and point-cloud tesselation to respectively reconstruct and quantify nanoscale organization of endogenous AMPAR in tissue. **b,** Representative 2D (left) and 3D (right) DNA-PAINT reconstruction of AP-GluA2, with projected GFP morphologic signal (10 planes) on the same FOV. (N_2D-frames_ = 20,000, N_3D-frames_ = 40,000). **c,** Zoom-in regions of **b** and corresponding reconstructed clusters and nanodomains after point-cloud tesselation analysis. Black arrows show the different representation of cluster/nanodmain object between 2D and 3D. **d,** Localization density map comparison of the AMPAR clusters pinned in **c**. **e,** Comparison of lateral and axial localization uncerntainty median and detection density (normalized at 100 frames) within the eGFP-defined area between 2D and 3D DNA-PAINT acquisition AP-GluA2. (N_2D_ = 11, N_3D_ = 12, *P ≤ 0.05, Mann-Whitney test). **f,** Surface object representation of localization pinned in **c** after point-cloud tesselation. **g,** Comparison of averaged projected axis measurement (2 axis in 2D, 3 axis in 3D) for each reconstructed cluster and nanodomains between 2D and 3D DNA-PAINT acquisitions of AP-GluA2. (N_2D-cluster_ = 550, N_3D-cluster_ = 719, N_2D-nd_ = 1573, N_3D-nd_ = 587, ****P ≤ 0.0001, Mann-Whitney test). **h,** Comparison of cluster and nanodomain volumes in 3D-DNA-PAINT acquisitions of AP-GluA2. **i.** Relative frequency of clusters containing different numbers of nanodomains in 2D and 3D DNA-PAINT acquisitions of AP-GluA2. (N_2D_ = 11, N_3D_ = 12). **j,** 3D rendering of 3D-DNA-PAINT acqusition in **b**. Insets shows MIPs XY, XZ and YZ of one AMPAR nanocluster. **k,** Correlation between spine head volume and AMPAR cluster area per spine, combining the 3D dendritic spine segmentation-illustrated with 3D rendering of **j**, with quantitative SMLM analysis. Lines indicate linear regressions with Pearson correlation coefficients (r) (N_2D_ = 196, N_3D_ = 199, ***P ≤ 0.001, ****P ≤ 0.0001, Pearson correlation). **Scale bar,** 2 μm (**b**), 1 μm (**j, k**), 500 nm (**c**), 100 nm (**j-inset**).

We first assessed how axial information influences the interpretation of endogenous AMPAR organization by comparing sequential 2D and 3D DNA-PAINT acquisitions of the same FOV (Fig. 3b). While both approaches recovered the overall receptor distribution, 3D localization additionally resolved the axial organization of receptor assemblies, particularly in densely labelled regions where spatially distinct structures overlapped in 2D. Localization datasets were subsequently analyzed using PoCA^39^, a point-cloud analysis framework based on Voronoi tessellation that performs multilevel segmentation directly in either 2D or 3D. Incorporating axial information altered both cluster boundaries and their internal nanodomain organization: structures appearing as single contiguous assemblies in 2D were separated into distinct clusters in 3D, while others revealed a more complex internal organization (Fig. 3c). Because point-cloud segmentation also depends on the spatial density of localizations, we further examined localization-density maps, which revealed differences between the 2D and 3D acquisitions (Fig. 3d). These observations prompted us to characterize localization performance and detection density across independent datasets before quantitatively comparing receptor nano-organization.

For each FOV, the neuronal morphology was defined from the diffraction-limited GFP channel using a manually generated region of interest (ROI). Only localizations and the resulting clusters and nanodomains within this region were retained for quantitative analysis, thereby excluding background detections. We then compared localization performance across independent 2D and 3D acquisitions. As expected, axial encoding increased the median lateral localization uncertainty from 8.79 nm in 2D to 18.59 nm in 3D, while providing a median axial localization uncertainty of 146.4 nm. Despite this expected trade-off in lateral precision, 3D acquisitions showed a higher detection density, increasing from 3.72 to 4.82 detections µm⁻² per 100 frames compared with 2D acquisitions (Fig. 3e). The achieved lateral localization uncertainty remained below the characteristic 70–100 nm dimensions of AMPAR nanodomains, while axial localization provided additional depth information to distinguish receptor assemblies positioned at different z coordinates within the imaging volume.

Having characterized localization performance across acquisition modes, we next assessed how incorporating axial information affected the quantitative description of AMPAR nano-organization. PoCA segmentation generated individual cluster and nanodomain objects directly from the 2D and 3D point clouds (Fig. 3f). Population-level analysis revealed differences in the measured dimensions of these objects, with cluster size increasing from 152.1 nm in 2D to 176.5 nm in 3D and nanodomain size from 34.35 nm to 68.03 nm, respectively (Fig. 3g). Notably, the characteristic nanodomain dimension obtained from the 3D analysis was within the ∼70–100 nm range previously reported for AMPAR nanodomains using independent super-resolution approaches^40^, although absolute size estimates depend on the segmentation and dimensionality of the analysis. 3D segmentation additionally captured the spatial distribution of receptor clusters and nanodomains across all three dimensions, enabling their organization to be better resolved over the 3D structure of dendritic spines (Fig. 3h). Finally, incorporating the axial dimension altered the distribution of nanodomains associated with individual clusters, revealing differences in their internal organization compared with 2D analysis (Fig. 3i).

Having established the quantitative differences introduced by 3D analysis, we next considered the spatial organization of endogenous AMPAR clusters within their cellular context. 3D rendering of the DNA-PAINT point cloud visualized individual receptor assemblies across all three spatial dimensions, providing the basis for relating their nanoscale organization to the dendritic spines in which they are localized (Fig. 3j). To quantitatively establish this relationship, we integrated the single-molecule point clouds with volumetric morphological segmentation using RESPAN^41^, a deep learning-based framework for neuronal morphology analysis. RESPAN was applied to the deconvolved diffraction-limited channel to segment dendritic shafts and individual spines in 3D (Fig. 3k). The resulting morphological segmentation was registered with the corresponding DNA-PAINT reconstruction, enabling receptor clusters and nanodomains to be assigned to individual spines and quantitatively related to their morphology. We then correlated spine head volume with the total AMPAR cluster area associated with each spine (Fig. 3k). A significant positive correlation was observed for both 2D (r = 0.273, P < 0.001) and 3D (r = 0.318, P < 0.0001) SMLM analyses, consistent with previous observations linking dendritic spine size to AMPAR-mediated synaptic responses^42^. Together, these results demonstrate that ALPINE bridges molecular and cellular scales, enabling the 3D nanoscale organization of endogenous proteins to be quantitatively resolved within the architecture of intact tissue.

### ALPINE for live molecular dynamics of endogenous proteins in tissue

Molecular mobility is a key determinant of protein function, particularly at synapses where the dynamic exchange, confinement and stabilization of membrane proteins contribute to synaptic transmission and plasticity. SPT studies in cultured neurons have established that postsynaptic membrane proteins exhibit distinct dynamic behaviours according to their molecular interactions and local synaptic environment, with AMPARs providing a well-characterized example of dynamic exchange between synaptic and extrasynaptic membrane domains^7,43,44^. Extending these measurements to organized brain tissue requires maintaining single-molecule sensitivity during prolonged live imaging while preserving the cellular environment in which these dynamics occur. Building on the optical and labelling developments described above, ALPINE enables live SPT of endogenous proteins in tissue, combining the high temporal resolution and gentle illumination of LLSM with cell-specific endogenous labelling.

To validate the ability of ALPINE to distinguish protein-specific molecular dynamics, we extended the AP-tag/eSA labelling strategy from AP-GluA2 to a second established knock-in model, AP-NLGN1^45^, and tracked both endogenous proteins in organotypic brain slices using AIO-LLSM (Fig. 4a). These proteins were selected to represent distinct modes of postsynaptic membrane dynamics: AMPARs undergo lateral diffusion and dynamic exchange between synaptic and extrasynaptic membrane domains^7,43^, whereas NLGN1 is a trans-synaptic adhesion protein exhibiting stronger diffusional confinement at synapses through interactions with presynaptic adhesion partners and postsynaptic scaffolding proteins^45^. Their distinct synaptic roles and organization therefore provide a biological reference for assessing whether ALPINE can resolve expected differences in endogenous protein trafficking in tissue. Individual detections were subsequently linked into trajectories using TrackPy^46^, and their mobility was characterized from the apparent diffusion coefficient (D) and anomalous diffusion exponent (α) derived from mean-squared displacement (MSD) analysis.

**Fig. 4.**
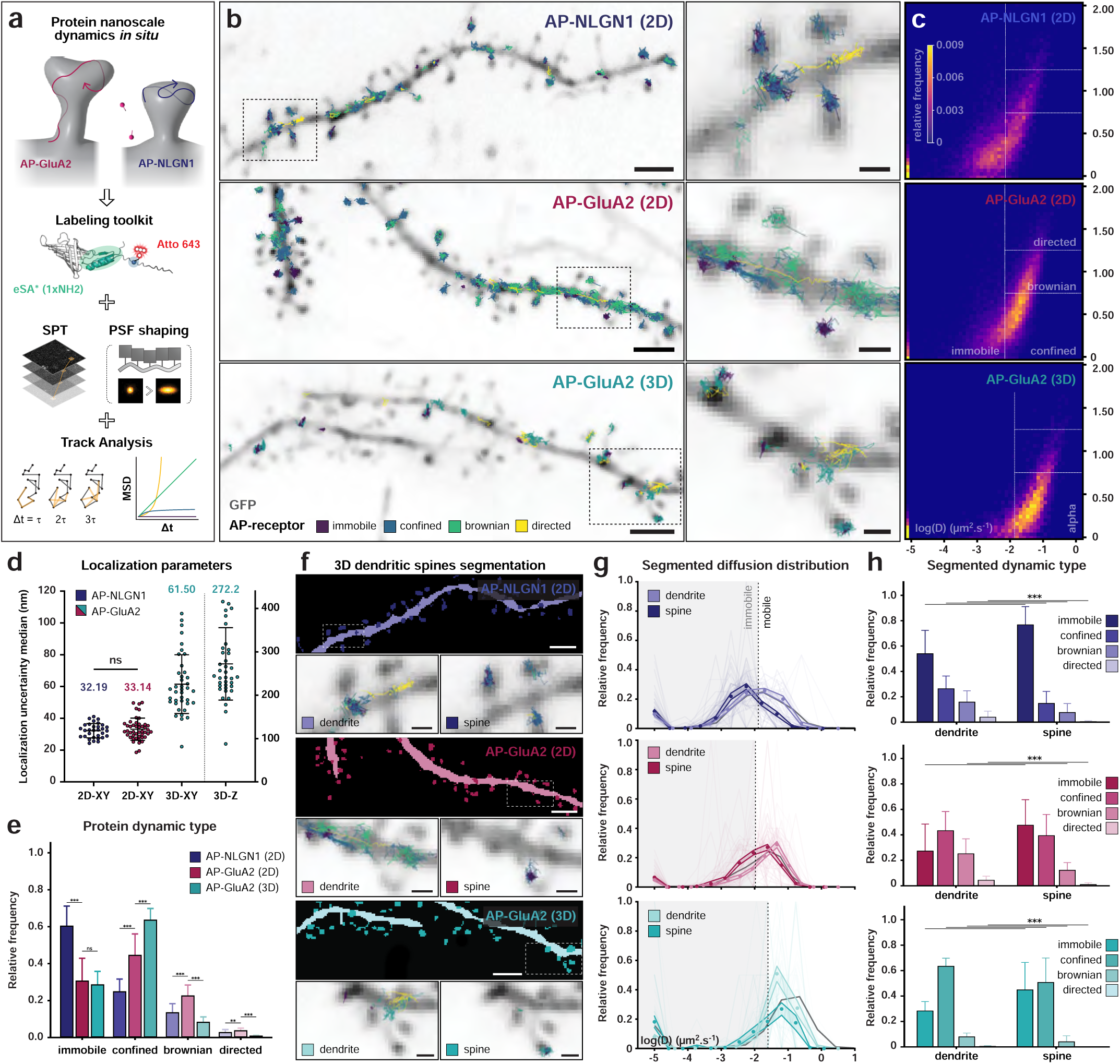
SPT-AO-LLSM and biotin-avidin labelling toolkit to visualize *in situ* protein nanoscale dynamics in living brain tissue. **a,** Schematic representation of the imaging pipeline using biotin-avidin strategy adapted for SPT imaging and track dynamics analysis to respectively reconstruct and quantify nanoscale dynamics of endogenous NLGN1 and AMPAR in living tissue. **b,** Representative dynamics of AP-NLGN1 (top) and AP-GluA2 receptors in 2D (middle) and 3D (bottom) with projected GFP morphologic signal (10 frames). MSD analysis allows to classify receptors dynamics (color-coded) from immobile to directed movements. Insets (right) zoom-in highlighted left areas. **c,** Tracks relative frequency depending on the diffusion coefficient (log(D)) and the anomalous diffusion exponent (alpha) for respetive receptors and acquisition type in **b**. White dotted line delineate the different dynamic type. **d,** Comparison of lateral and axial localization uncerntainty median per acquisition within the eGFP-defined area for AP-NLGN1, 2D-AP-GluA2 and 3D-AP-GluA2. (N_AP-NLGN1_ = 33, N_2D-AP-GluA2_ = 48, N_3D-AP-GluA2_ = 41, one-way ANOVA test). **e,** Relative frequency of protein dynamic type for corresponding acquisition in **d**. (N_track-AP-NLGN1_ = 14,328, N_track-2D-AP-GluA2_ = 21,340, N_track-3D-AP-GluA2_ = 7,019, **P ≤ 0.01, ***P ≤ 0.001, weighted t-test). **f,** 3D dendritic spines segmentation masks for zoom-in area in **b**. Insets-highlighted on mask’s representation-show tracks compartmentalisation between spine and dendrite. **g,** Relative distribution comparison of segmented dynamic diffusions for each condition in **f**. **h,** Relative frequency of segmented protein dynamic type for corresponding acquisition in **d**. (N_track-AP-NLGN1-dendrite_ = 1,823, N_track-2D-AP-GluA2-dendrite_ = 2,525, N_track-3D-AP-GluA2-dendrite_ = 470, N_track-AP-NLGN1-spine_ = 2,013, N_track-2D-AP-GluA2-spine_ = 2,346, N_track-3D-AP-GluA2-spine_ = 222, **P ≤ 0.01, ***P ≤ 0.001, weighted t-test). **Scale bar,** 2 μm (**b, f-mask**), 500 nm (**b-inset, f-inset**).

We next compared protein dynamics across three acquisition conditions: AP-NLGN1 in 2D, AP-GluA2 in 2D and 3D (Fig. 4b). Based on their D and α values, trajectories were assigned to dynamic regimes ranging from directed to confined motion using previously established criteria^47,48^, with an additional immobile class comprising trajectories with no detectable displacement or motion below the resolvable range. Representative trajectory maps revealed distinct spatial distributions, with NLGN1 trajectories predominantly associated with dendritic spines, whereas AP-GluA2 trajectories were more broadly distributed along dendritic shafts and spines. In addition, 3D-SPT of AP-GluA2 recovered axial components of receptor motion that are absent from 2D trajectories (Fig. 4b). Mapping the trajectories in the log(D)-α parameter space further revealed distinct distributions across the three acquisition conditions, reflecting differences in their dynamic behaviours (Fig. 4c).

We next quantified localization performance across the three conditions before comparing their dynamic classifications. Lateral localization uncertainty was similar for 2D AP-NLGN1 and AP-GluA2 acquisitions, with median values of 32.19 and 33.14 nm, respectively, whereas 3D AP-GluA2 imaging provided axial localization with a median uncertainty of 272.2 nm at the cost of an approximately twofold increase in lateral uncertainty to 61.50 nm (Fig. 4d). These acquisition modes therefore provide complementary capabilities: 2D SPT offers higher lateral localization precision, whereas 3D SPT incorporates axial information that can facilitate the assignment of molecular trajectories across the 3D architecture of cellular structures such as dendritic spines. Despite these differences in localization performance, trajectory classification recovered the expected protein-specific dynamics (Fig. 4e). Compared with GluA2, NLGN1 showed a greater proportion of restricted trajectories, consistent with its stronger stabilization through trans-synaptic adhesion and postsynaptic scaffold interactions, whereas the larger mobile fraction of AMPARs is consistent with its dynamic exchange and lateral diffusion between synaptic and extrasynaptic membrane domains^43,49^. Together, these complementary acquisition modes provide flexibility to prioritize localization precision or 3D spatial information according to the biological question, while enabling quantitative comparison of molecular dynamics across experimental conditions.

Having established protein- and dimensionality-dependent differences in molecular dynamics, we next asked how these behaviours relate to the local neuronal morphology. To add this cellular context to the SPT analysis, RESPAN segmentation^41^ was used to define dendritic shafts and individual spines, and the resulting 3D masks were spatially registered with the corresponding single-molecule trajectories (Fig. 4f). This correlation enabled individual tracks to be assigned to dendritic or spine compartments, providing a framework to further investigate receptor dynamics according to their local cellular environment.

Compartment-specific analysis revealed distinct mobility distributions between dendritic shafts and spines across the three conditions (Fig. 4g). For both NLGN1 and GluA2, trajectories within spines were shifted toward lower apparent diffusion coefficients compared with those assigned to dendritic shafts, indicating reduced molecular mobility within the spine compartment. This compartment-dependent restriction was also observed for GluA2 in 3D, showing that differences in receptor mobility between dendritic shafts and spines are retained when axial motion is incorporated into the analysis.

Consistent with these differences in apparent diffusion, classification of trajectories into dynamic states revealed a compartment-dependent redistribution of molecular behaviours (Fig. 4h). For both NLGN1 and GluA2, spines contained a greater proportion of restricted trajectories than dendritic shafts, with an increase in immobile and confined populations and a corresponding reduction in more mobile states. This redistribution is consistent with the increased stabilization of postsynaptic membrane proteins within spines, through receptor trapping and interactions with synaptic scaffolds. Similar compartment-dependent differences were observed for 3D GluA2 trajectories, demonstrating that ALPINE can resolve protein-specific and compartment-dependent molecular dynamics while relating individual trajectories to the 3D cellular architecture of intact tissue.

### ALPINE links single-molecule organization and dynamics to cellular context

To illustrate the analysis possibilities enabled by ALPINE, we combined single-molecule quantification with advanced 3D morphological analysis. This integration enables molecular organization and dynamics to be related to defined cellular features, providing a framework to investigate how local morphology is associated with protein organization and behaviour. We applied this approach to dendritic spines, combining quantitative spine morphology with nano-organization and trafficking measurements of endogenous AMPARs presented in Figures 3 and 4.

Dendritic spine morphology was reconstructed from the diffraction-limited GFP channel following deconvolution (Fig. 5a) and segmented in 3D using RESPAN^41^. This automated segmentation identifies dendritic shafts and individual spines, enabling their visualization within the complete dendritic morphology (Fig. 5b). Dendritic spines exhibit diverse morphologies, commonly described as filopodia, stubby, thin and mushroom spines, which are associated with differences in spine maturation, stability and synaptic function^50^. Representative examples illustrate the initial 3D GFP volume, segmentation mask and corresponding surface representation for each morphology (Fig. 5c). To characterize this morphological diversity without predefined spine categories, 26 morphological parameters describing the entire spine, spine head and spine neck were extracted from RESPAN and analysed using principal component analysis (PCA) followed by K-means clustering (Fig. 5d). The resulting clusters were subsequently associated with the four commonly described spine morphologies based on their characteristics and visualized using three selected parameters (head cross-section, neck cross-section and spine length) to illustrate the differences between categories (Fig. 5d).

**Fig. 5.**
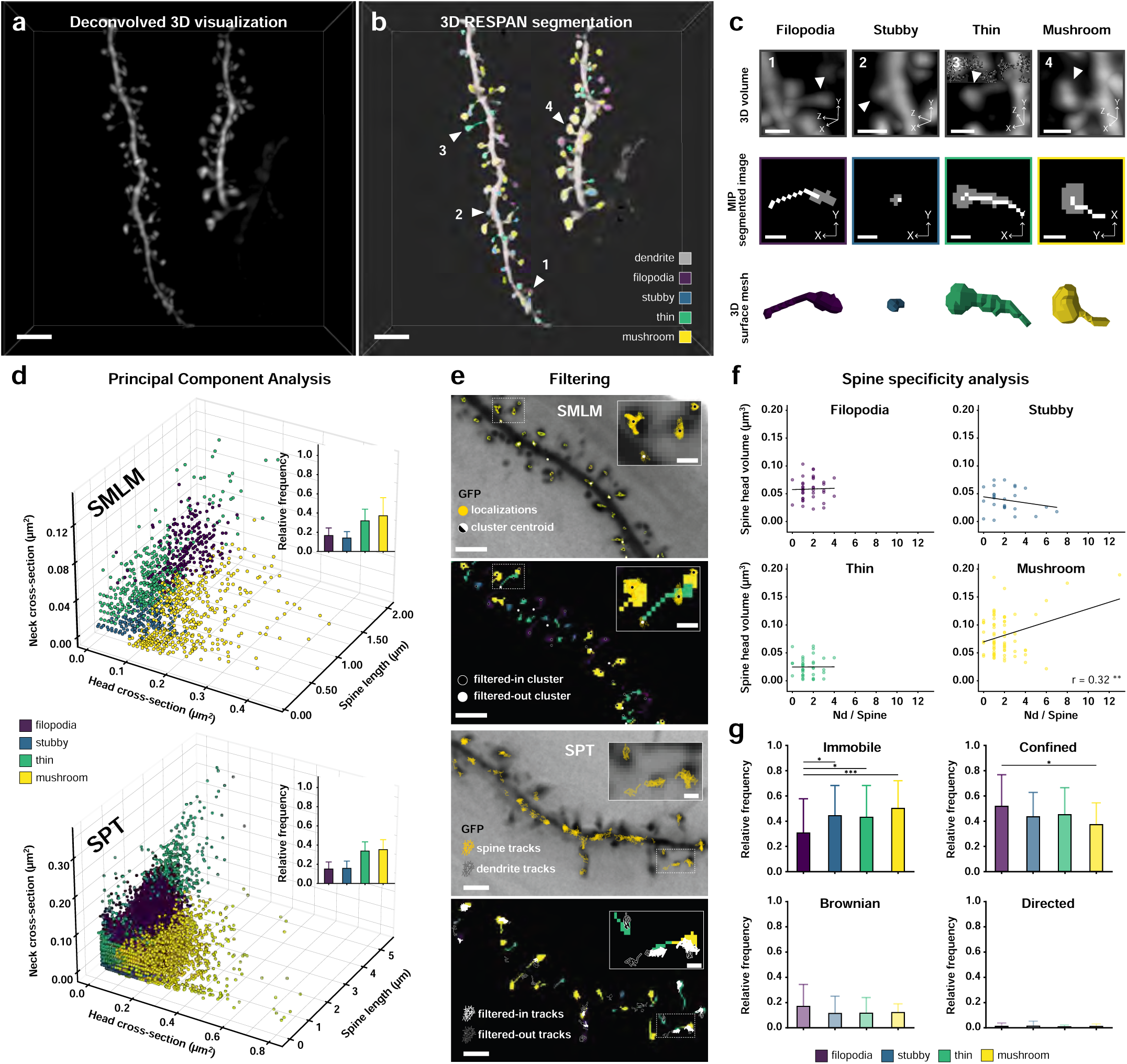
Correlation of single-molecule measurements with 3D spine morphology. **a,** Representative deconvolved 3D visualization of dendritic morphology from the GFP channel. **b,** Corresponding 3D RESPAN segmentation of the dendritic shaft and individual spines, color-coded according to spine morphology. **c,** Representative filopodia, stubby, thin and mushroom spines, highlighted by arrowheads 1-4 in **b**, showing the initial 3D GFP volume (top), corresponding MIP of the segmentation mask (middle) and 3D surface representation (bottom). **d,** PCA-based morphological classification of individual spines from SMLM (top) and SPT (bottom) datasets. Individual spines are color-coded according to their morphological classification. Insets show the relative frequency of each spine type (N_spines-SMLM_ = 1,772, N_spines-SPT_ = 25,526). **e,** Representative spatial filtering of SMLM clusters (top) and SPT trajectories (bottom) according to the corresponding 3D RESPAN segmentation masks. **f,** Correlation between spine-head volume and the number of AMPAR nanodomains per spine for filopodia, stubby, thin and mushroom spines. Lines indicate linear regressions with Pearson correlation coefficients (r) (N_filopodia_ = 37, N_stubby_ = 26, N_thin_ = 37, N_mushroom_ = 71, **P ≤ 0.01, Pearson correlation). **g,** Relative frequency of immobile, confined, Brownian and directed AMPAR trajectories according to spine morphology (N_immobile-filopodia_ = 62, N_immobile-stubby_ = 75, N_immobile-thin_ = 147, N_immobile-mushroom_ = 768, N_confined-filopodia_ = 104, N_confined-stubby_ = 73, N_confined-thin_ = 153, N_confined-mushroom_ = 581, N_brownian-filopodia_ = 34, N_brownian-stubby_ = 19, N_brownian-thin_ = 39, N_brownian-mushroom_ = 188, N_directed-filopodia_ = 2, N_directed-stubby_ = 2, N_directed-thin_ = 2, N_directed-mushroom_ = 15, *P ≤ 0.05, ***P ≤ 0.001, weighted t-test. **Scale bar,** 3 μm (**a, b**), 2 μm (**e**),1 μm (**c-top**), 500 nm (**c-middle, e-insets**).

We next used these morphological segmentations as a spatial reference for the single-molecule measurements. Registration of the diffraction-limited morphology with the corresponding SMLM or SPT datasets enabled molecular measurements to be assigned to individual spines (Fig. 5e). To account for the spatial uncertainty between single-molecule localizations and the diffraction-limited segmentation, the morphological masks were expanded according to the localization uncertainty of each dataset. For SMLM, clusters were assigned to spines when their centroid fell within the corresponding expanded mask, retaining 50.6 % of clusters. For SPT, trajectories were assigned based on their occupancy, retaining tracks with ≥95% of localizations within a single spine or dendritic compartment (22.0%). Together, these approaches enabled nanoscale AMPAR organization and mobility to be quantitatively related to defined spines.

We first examined whether AMPAR nano-organization varied with spine morphology by quantifying the number of nanodomains associated with individual spines. Pearson correlation analysis was used to assess the relationship between spine-head volume and nanodomain number for each morphological class (Fig. 5f). indicating that larger mushroom spine heads were associated with a greater number of AMPAR nanodomains, consistent with previous observations that larger synapses can accommodate multiple nanoscale modules of synaptic proteins^51^. Finally, we applied the same structural assignment to live SPT measurements to determine whether AMPAR mobility differed between spine morphologies. Individual trajectories were classified according to their diffusive behavior and assigned to filopodia, stubby, thin or mushroom spines. Comparison of the relative frequencies of immobile, confined, Brownian and directed trajectories revealed morphology-dependent differences in AMPAR mobility (Fig. 5g). The fraction of immobile trajectories was significantly lower in filopodia than in stubby, thin and mushroom spines, whereas confined trajectories were significantly more frequent in filopodia than in mushroom spines. Together, these analyses demonstrate that ALPINE can relate both nanoscale molecular organization and dynamics to defined 3D cellular structures, providing a framework to investigate how local cellular architecture is associated with endogenous protein organization and behaviours in intact tissue.

## Discussion

Here, we establish ALPINE as a multimodal framework that brings quantitative imaging of endogenous molecular organization and dynamics into complex biological tissue. By combining AO-LLSM for 3D single-molecule imaging within optically complex samples, endogenous and tuneable molecular labelling, and dedicated analysis workflows to link molecular events to their native context, ALPINE integrates nanoscale molecular organization and dynamics with cellular context into a modulable framework. Using endogenous cell-surface proteins as models, we demonstrate 3D-SMLM and 3D-SPT in fixed and living brain tissue, while correlating molecular behaviours with their morphological context through neuronal compartments and classified spine types. Rather than solely resolving single molecules within tissue, this integration enables their endogenous organization and dynamics to be quantitatively examined within defined 3D cellular environments.

Previous developments have established the potential of bringing SMLM from cultured cells into preserved tissue. REALM^52^ and DL-AO^53^ demonstrated that AO-SMLM could resolve nanoscale structures in simulated or real brain sections, respectively, illustrating the potential of accessing molecular-scale information within a physiologically relevant context. ALPINE moves into live endogenous molecular dynamics in tissue, combining 3D single-molecule localization and tracking with the cellular context in which these molecular behaviours occur. Previous studies have established receptor tracking in brain slices and 3D-SPT in organotypic brain tissue using nanoparticle-based probes^10,54^, while 3D tracking of endogenous AMPARs has been demonstrated in cultured neurons^55^. More broadly, the ability to follow individual endogenous proteins in living tissue creates opportunities to investigate molecular dynamics directly within their native cellular environment. Applied here to AMPARs and NLGN1, this capability enables protein-specific dynamics to be compared across dendritic and synaptic compartments and related to local spine morphology, providing a route to investigate how synaptic architecture and endogenous molecular behaviour shape synaptic function.

By combining AO with LLSM, ALPINE corrects tissue-induced aberrations without requiring guide stars or wavefront sensing during imaging, restoring single-molecule detection performance at depth, as quantified by detection number and localization uncertainty. By reducing optical aberration-induced biases, AO provides more consistent imaging conditions across tissue depths, improving the reliability and comparability of quantitative single-molecule measurements. This capability could extend molecular measurements across different depths and cellular environments within the same tissue, providing access to biological variability that would otherwise remain difficult to distinguish from depth-dependent optical degradation. As previously proposed for thick neuronal samples^56^, ALPINE combines light-sheet illumination with programmable PSF engineering through the DM for 3D single-molecule localization. Unlike conventional cylindrical-lens approaches, the DM introduces controlled Zernike astigmatism directly onto the aberration-corrected wavefront, without introducing additional field distortions or magnification changes^52^. This approach preserves a compact PSF and efficient photon collection, both critical for accurate 3D single-molecule localization. For SMLM, 3D analysis altered the measured nanoscale organization of AMPARs compared with lateral projections, with nanodomains that overlapped in 2D being resolved as spatially distinct structures along the axial dimension. Similarly, incorporating axial displacement into SPT modified the measured mobility of a subset of trajectories, with tracks appearing confined or immobile from their lateral displacement alone exhibiting substantial motion along the axial dimension. Together, these capabilities address two complementary sources of ambiguity in tissue single-molecule imaging: AO reduces aberration-induced measurement biases to support reproducible quantification across heterogeneous imaging depths, while 3D localization provides a more complete representation of molecular organization and motion within native cellular environments.

Accessing endogenous molecular behaviour in tissue also requires labelling strategies that combine molecular specificity with the signal intensity and flexibility required for single-molecule imaging modalities. This is particularly important for quantitative measurements of molecular organization and dynamics, where exogenous expression can alter protein abundance and potentially perturb the distribution or interactions of the molecules being measured. To meet these requirements, we developed eSA, a new streptavidin-based labelling probe optimized for single-molecule imaging of endogenous AP-tagged proteins in tissue. Building on the AP-tag/biotinylation strategy for endogenous GluA2^25^, we developed an engineered streptavidin-based labelling toolkit that increases labelling signal while maintaining target specificity, meeting the requirements of single-molecule imaging of endogenous surface receptors. Importantly, the introduction of secondary labelling sites provides a modular interface through which the final imaging probe can be adapted while preserving specific recognition of the endogenous target. This enables signal amplification and flexible reporter functionalization, as demonstrated here by coupling DNA strands through ALFA tag nanobodies for DNA-PAINT. Such a modular architecture could facilitate adaptation to different imaging modalities and multiplexed imaging of distinct molecular targets through orthogonal labelling strategies, without requiring redesign of the primary target-recognition system.

To integrate nanoscale molecular measurements with their native cellular context, we incorporated an existing deep-learning, nnU-Net-based approach for automated 3D segmentation of neuronal morphology into the ALPINE analysis workflow^41^. The available model enabled identification of dendritic compartments and individual spines from our diffraction-limited LLSM acquisitions, allowing receptor clusters and trajectories to be assigned to their local 3D morphological environment. Combined with spine morphological classification, we observed a positive correlation between spine-head volume and AMPAR nanodomain number specifically in mushroom spines, while SPT revealed morphology-dependent differences in receptor mobility, with filopodia displaying a lower fraction of immobile trajectories than other spine classes and a higher fraction of confined trajectories than mushroom spines. These results demonstrate how molecular organization and dynamics can vary according to the local morphological environment in which they are measured. More broadly, integrating single-molecule measurements with automated morphological analysis provides a framework for investigating such molecular heterogeneity across structurally diverse cellular compartments.

Several aspects of the current implementation define opportunities for further development. First, aberration correction is estimated from the diffraction-limited morphological channel and transferred to the single-molecule channel. Direct wavefront optimization from single-molecule emission could eliminate this inter-channel transfer and provide correction tailored to the molecules being measured, as emerging approaches for single-molecule-based aberration estimation suggest^53^. Second, the astigmatic PSF used here provides a simple approach for 3D localization but retains substantially lower axial than lateral localization precision. More advanced PSF engineering, including extended-range Tetrapod PSFs, combined with recent learning-based localization approaches could improve the accuracy and axial range of 3D molecular measurements, including under aberrated conditions^57^. Further improvements in cross-channel registration could enable molecular measurements to be related to cellular morphology at finer spatial scales, for example by resolving cluster positions within individual spines or the local membrane position of trajectories. The demonstrated labelling strategy is currently restricted to extracellularly accessible, AP-tagged proteins, and extension to intracellular targets will require alternative labelling chemistries compatible with endogenous single-molecule imaging. Finally, while AO compensates wavefront distortions, scattering ultimately limits imaging performance and accessible depth. Combining ALPINE with clearing approaches compatible with fixed or living tissue, including recently developed minimally invasive clearing media^58^, could therefore further expand the accessible imaging volume.

Despite these current limitations, the modular architecture of ALPINE should facilitate its adaptation to different molecular targets and biological samples. AIO could operate from alternative structural markers through the selection of an appropriate image-quality metric, while the imaging framework could be combined with labelling strategies providing sufficient specificity and signal for single-molecule detection. Similarly, where suitable segmentation approaches are available, the registration and molecular-assignment workflow could associate molecular measurements with other cellular structures and morphological features. Together, these components provide a configurable framework for adapting molecular measurements to the biological system under investigation.

Together, these current capabilities and opportunities for further improvement establish ALPINE as a framework for studying molecular behaviour directly within complex biological environments. Importantly, the present implementation already demonstrates that combining AO and optical sectioning, adaptable endogenous labelling, and dedicated analysis provides access to both the nanoscale organization and dynamics of molecular populations within defined cellular structures in tissue. This is particularly relevant at synapses, where molecular composition, morphology and function are highly heterogeneous, yet how this diversity shapes the nanoscale organization and mobility of endogenous receptors remains largely unresolved^1^. Relating endogenous molecular behaviour to synapse morphology, identity, activity or plasticity could therefore reveal how nanoscale molecular mechanisms are specialized across distinct synaptic environments.

The need to resolve molecular behaviour within its local cellular environment extends well beyond the synapse, across biological systems in which spatially specialized cellular domains determine molecular function. At the axon initial segment, the nanoscale organization and remodeling of ion channels and scaffolding proteins are tightly linked to neuronal excitability^59^. Similarly, in polarized epithelia, organoid models enable molecular dynamics to be examined across functionally distinct apical, lateral and basal membrane domains, as demonstrated for clathrin-mediated endocytosis in intestinal epithelial organoids^60^. In plants, specialized membrane domains such as plasmodesmata illustrate how nanoscale molecular organization and dynamics regulate trafficking and signaling between neighboring cells^61^, while recent LLSM-STORM imaging of receptor-like kinases in intact leaf tissue demonstrates the feasibility of resolving such molecular organization directly within plant tissue^62^. These diverse biological systems share a common challenge: understanding how molecular organization and dynamics are shaped by the local cellular architecture and environment in which they operate. By connecting single-molecule measurements with this biological context, ALPINE provides a route from observing nanoscale molecular behaviour to investigating how it varies across cells, compartments and functional states in complex biological systems.

## Methods

### Animals

Three knockin (KI) mouse lines were used in this study: the Thy1-YFP-H mouse strain (JAX stock #003782)^1^, the AP-GluA2 KI mouse line, generated as described previously^2^, and the AP-NLGN1 KI mouse line, generated as described previously^3^. The mouse strains used in this study were bred in the EOPS (Free from Specific Pathogens) facility of the common animal facility of Bordeaux and maintained and manipulated at the Platform In vivo (PIV-EXPE) facility of the Bordeaux Neurocampus. Animals were housed in 12/12 light/dark with ad libitum feeding. Every effort was made to minimize the number of animals used and their suffering. The experimental designs and all procedures were in accordance with the European guide for the care and use of laboratory animals and the animal care guidelines issued by the animal experimental committee of Bordeaux Universities (CE50; animal facilities authorizations B33-063-941 and B33-063-940 respectively; ethical project authorizations #30029 and #38390).

### Organotypic hippocampal slice preparation

Organotypic hippocampal slices were prepared and cultured using the interface method^4,5^, as previously described in detail^2,6^. Briefly, at postnatal day 5 -8 animals of either sex were anesthetized on ice and decapitated. Hippocampi were dissected in ice-cold dissection medium composed of (in mM): 10 D-glucose, 4 KCl, 26 NaHCO_3_, 233.17 sucrose, 5 MgCl_2_, 1 CaCl^2^, 0.1% phenol red equilibrated with carbogen (95% O_2_, 5% CO_2_). Transverse 300 μm slices were cut with a tissue chopper (McIlwain), and transferred into ice-cold culture medium composed of: MEM powder (Sigma-Aldrich, #M4642), 20% inactivated horse serum (Gibco, #26050088), 1 mM L-glutamine (Thermo Fisher, #25030081), 1 mM CaCl_2_, 2 mM MgSO_4_, 1 mg/L insulin (Sigma, #I0516), 0.25 mM sodium L-ascorbate, 30 mM HEPES, 13 mM D-glucose, 5.2 mM NaHCO 3 (pH 7.3, 320 mOsm/L). Slices were then plated on 4 mm membrane segments (Millipore, #FHLC01300) placed on culture inserts (Millipore, #PICM0RG50) in 6-well plates containing 1 ml slice culture medium per well. Slices were maintained at 35 ◦ C with 5% CO 2 and the culture medium was replaced every 2-3 days.

### AP-GluA2 Organotypic hippocampal slice transduction

Slices were transduced^6^ with hSyn-BirA^ER^-IRES-Cre + hSyn-DIO-eGFP AAV9^2^ at DIV 1 by microinjection of the viruses (three to five pulses, 30 ms, 69 kPa) into the CA1 pyramidal cell layer with glass microelectrodes (∼1 to 2 megaohms; Science Products, GB150F-10P). BirA^ER^-IRES-Cre + DIO-eGFP were used at a dilution of 1:5000 to 1:20,000 in 1× phosphate-buffered saline (PBS). Viral titres ranged between 2.3-6.7 x 10^13^ GCP/mL. Upon returning the slices to the incubator the culture medium was supplemented with 10 μM D-biotin (for mSA labelling experiments) until experiments were performed.

### Protein production and purification

The momoneric streptavidin variants and the anti-ALFA tag nanobody (nbALFA*, passivated nanobody with a single lysine for coupling) were obtained as previously described. Briefly, BL21 CodonPlus (DE3)-RIPL cell pellets expressing mSA/eSA variants or nbALFA* were thawed, resuspended in TA 1X 5 mM Imidazole buffer supplemented with protease inhibitors and lysozyme, and sonicated. For mSA/eSA variants, Triton X-100 (1% final) was added. mSA/eSA variants were purified from inclusion bodies by washing the pellet three times with TA 1X 5 mM Imidazole, dissolving in 8M Urea, and refolding at 4°C by dropwise addition into PBS 1X. nbALFA* was purified from the soluble fraction. Both proteins were bound to Nickel-NTA Agarose gel (overnight for mSA/eSA variants, 2 h for nbALFA*), washed, and eluted with TA 1X 500 mM Imidazole. For nbALFA*, the eluate was dialyzed against PBS 1X at 4°C with NEDP1 protease, followed by a reverse His-tag affinity step to remove NEDP1 and MBP. Both proteins were further purified by size-exclusion chromatography (SEC, Superdex 75) on a FPLC system and concentrated using Amicon Ultra 15 tubes (10 kDa cutoff). Proteins were aliquoted in liquid nitrogen and stored at -80°C.

### Probe functionalization

Dye conjugation was performed at pH 8 (adjusted with NaHCO₃ 1 M). mSA/eSA variants were labeled with 7 eq (mol/mol) dye and 500 mM 3-(Benzyl dimethylamonio)propanesulfonate for 2 h at RT. nbALFA* was labeled with 9 eq (mol/mol) dye overnight at 4°C. Labeled proteins were desalted on PD MiniTrap G-25 columns equilibrated with PBS 1X, and the eluate was collected in 1 mL PBS 1X. Proteins were aliquoted in liquid nitrogen and stored at -80°C.

### bAP-GluA2 labelling

eSA labeling experiments were performed on AP-GluA2 or AP-NlG1 organotypic slices from 12 to 15 DIV. Samples were washed twice for 10 min in Tyrode’s solution containing 10 mM d-glucose, 10 mM Hepes, 120 mM NaCl, 3.5 mM KCl, 2 mM MgCl2, 2 mM CaCl2, 2 mM NaHCO3, and 1 mM Na-pyruvate (pH 7.3; 300 mOsm) with 1% biotin-free BSA. For the development of the biotin–avidin labeling toolkit and AMPAR nanoscale organization imaging by DNA-PAINT, slices were incubated for 20 min in approximately 30 µl of Tyrode-BSA containing 200 nM eSA. For DNA-PAINT experiments, the eSA-4×ALFA variant was used, followed by labeling with an anti-ALFA single-domain antibody conjugated to the F3 DNA docking strand. Samples were then washed twice for 10 min in Tyrode-BSA before fixation (see below). For live single-particle tracking (SPT) experiments, slices were incubated for 5–10 min in Tyrode-BSA containing 4–5 nM fluorescent dye-conjugated eSA* (1×NH_2_), to minimize AMPAR internalization. Samples were then imaged directly without fixation.

### Slices labelling for DNA-PAINT experiments

Thy1-YFP organotypic slices were fixed at 15 DIV without treatment, while APGluA2 slices were fixed after eSA live staining as described above. Each slice was placed individually in a well of a 24-well plate and incubated for a few seconds with 800 µl of a pre-fixation solution consisting of Tyrode’s buffer containing 2% paraformaldehyde (PFA) and 2% sucrose. The pre-fixation solution was then removed, and slices were incubated with 800 µl of 4% PFA and 4% sucrose in PBS at 4 °C for 3 h. Slices were washed three times with PBS, then incubated for 30 min in PBS containing 50 mM ammonium chloride under gentle agitation to quench free aldehydes. After three additional PBS washes, permeabilization was performed 24 h at 4 °C in PBS containing 1% Triton X-100 under gentle agitation. Following three PBS washes, blocking was carried out for 2 h at room temperature with gentle agitation in a blocking solution composed of 3% IgG-free BSA, 1% sheared salmon sperm DNA (Thermo Fisher Scientific, AM9680), 1% Triton X-100, and 0.2% fish gelatin in PBS. Anti-GFP and/or anti-ALFA single-domain antibodies (MASSIVE-TAG-X2-FAST anti-GFP and MASSIVE-TAG-Q-FAST anti-ALFA; Massive Photonics GmbH), conjugated to FAST DNA-PAINT docking strand sequences (F1 and F3, respectively), were diluted 1:200 in antibody incubation solution containing 3% IgG-free BSA, 1% sheared salmon sperm DNA, 0.02% Tween-20, 1 mM EDTA, and 0.2% fish gelatin in PBS. Slices were transferred to individual wells of a 96-well plate containing 200 µl of antibody solution and incubated overnight at 4 °C under agitation. After labelling, slices were transferred back to the 24-well plate, washed three times with PBS, and post-fixed for 30 min at room temperature in PBS containing 2% PFA. Finally, slices were washed three times in PBS and stored in fresh PBS until imaging.

### AO-LLSM optical setup

Imaging was performed using a custom AO-LLSM system^7^ based on the original LLSM design^8^. Briefly, the excitation beam was shaped using a programmable spatial light modulator to generate an optical lattice, spatially filtered by an annular mask, and projected into the sample through a 20×, 0.60-NA water-immersion excitation objective (TL20X-MPL, Thorlabs). For all acquisitions, an annular mask with inner and outer NAs of 0.44 and 0.55, respectively, was used to generate the light sheet. The light-sheet position along the detection axis was controlled by a galvanometric mirror conjugated to the back focal plane of the excitation objective. Fluorescence was collected orthogonally to the excitation axis using a 25×, 1.1-NA water-immersion detection objective (Apo LWD 25×W, N25X-APO-MP, Nikon) and imaged onto an ORCA-Fusion BT sCMOS camera (C15440-20UP, Hamamatsu Photonics). The effective pixel size in the sample plane was 95 or 108 nm, as determined by spatial calibration of the imaging system. Excitation was provided by 488, 560 and 642 nm laser lines. The detection path incorporated a temperature-stabilized 52-actuators DM (Mirao 52-es, Imagine Optic) positioned in a plane conjugate to the back focal plane of the detection objective. The DM was used to compensate for static instrumental and sample-induced optical aberrations and, for 3D single-molecule imaging, to introduce a controlled astigmatic deformation of the detection PSF. A Shack–Hartmann WFS was incorporated into a separate calibration path and used for DM calibration and correction of static instrumental aberrations. The optical design, alignment procedure, DM calibration and correction of static aberrations have been described previously. Microscope operation and image acquisition were controlled using the original LLSM control software, whereas AO operations were managed by a custom LabVIEW application running in parallel with the microscope-control software. The AO software was developed in collaboration with Imagine Optic using its software development kit and provided independent control of the DM and implementation of the image-based optimization procedures. This architecture enabled aberration correction to be performed directly from images acquired by the microscope without modification of the primary LLSM control software.

### Active image optimization

Sample-induced aberrations were corrected locally using the previously developed active image optimization (AIO) procedure^7^. For each region selected for single-molecule imaging, AIO was performed on the GFP/YFP-labelled neuronal morphology within the same FOV. The procedure consisted of two sequential steps: correction of the axial defocus between the light sheet and the detection focal plane, followed by sensorless correction of aberrations in the detection path. First, the axial position of the light sheet was optimized by varying its position along the detection axis using the excitation-path galvanometric mirror. Five image stacks were acquired with different light-sheet axial offsets spanning −2 to +2 µm. For each offset, an image-based intensity metric was calculated from the maximum-intensity projection of the acquired stack, and the optimal light-sheet position was determined by Gaussian fitting. The resulting offset was then applied to the galvanometric mirror to maximize the overlap between the illumination plane and the focal plane of the detection objective. Detection-path aberrations were subsequently corrected using DM and the previously described 3N+ sensorless modal optimization algorithm^7^. Zernike modes were optimized sequentially by applying negative, null and positive amplitudes of each mode with the DM and measuring the corresponding image-quality metric. The metric was calculated from the spatial-frequency content of the GFP/YFP morphological signal by integrating the Fourier-domain energy between 500 and 2,000 mm⁻¹. For each Zernike mode, a second-order polynomial was fitted to the inverse of the measured metric to determine the coefficient maximizing image quality. When the optimum fell outside the initially sampled range, additional modal amplitudes were iteratively tested according to the 3N+ procedure. The correction comprised two successive optimization iterations. The first iteration used a modal sampling amplitude of 100 nm RMS, followed by a second iteration using 50 nm RMS to refine the correction. In each iteration, seven Zernike modes were sequentially optimized in the following order: primary spherical aberration 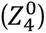, the two orthogonal astigmatism modes 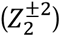, the two orthogonal coma modes 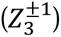, and the two orthogonal trefoil modes 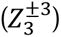. The final DM shape was retained for the subsequent single-molecule acquisition. Because sample-induced aberrations vary spatially within heterogeneous tissue, AIO was performed independently for each selected imaging region. Each AIO procedure takes less than 1 min. The GFP/YFP morphological channel therefore served as a structural guide signal for the optimization, allowing the aberration correction to be determined without using the single-molecule channel during the AIO procedure. After correction, imaging was switched to the single-molecule channel while maintaining the optimized DM shape.

### Image plane stabilization

We applied a two-step drift-correction method for DNA-PAINT imaging: periodic axial drift correction during acquisition, followed by post-processing lateral drift cancellation. To stabilize the axial (Z) position of the image plane, we acquired small stacks (typically 15 × 15 × 10 µm³) of the GFP channel (488 nm excitation) repeatedly, approximately every 2 minutes, interleaved with 1000–2000 DNA-PAINT frames (560 or 642 nm excitation) (Fig. S2a). Since fiducial markers could not be introduced into the brain slices, we used the GFP-channel signal to detect sample drift. This channel presents high-contrast structures such as bright dendrites or axons against a uniform low background. GFP stacks are acquired by scanning the light sheet and the detection objective simultaneously along the Z axis, i.e., along the detection objective’s optical axis. The 3D drift vector between stack S_n_ at time point t_n_ and stack S_0_ at t_0_ was computed from 2D cross-correlations of maximum-intensity projections (MIPs) along X, Y, and Z. To ensure robustness and reliability of the drift measurement, we normalized each MIP image by dividing pixel intensities by the total image intensity^9^. This normalization equalizes energy levels between images, mitigates biases such as photobleaching, and suppresses intensity offsets. Next, 2D cross-correlations are computed using conventional FFT-based algorithms. Subpixel precision of the drift vector is achieved by fitting the cross-correlations with a 2D Gaussian to localize the peak position. Although we measured the drift vector along X, Y, and Z, we stabilized only the Z position during long DNA-PAINT acquisitions by applying translations to the light sheet and the detection objective (Fig. S2a, bottom). We could not use the sample stages (Physic Instrument U-521 miniature linear stage) to correct lateral drift because their minimum step is 300 nm, which is too coarse for repositioning. In contrast, the positioning resolution of the light-sheet and the detection objective stages (Physik Instrument Piezo P621.1CD) is 3.75 nm and 0.4 nm, respectively. Following axial stabilization during acquisition, residual lateral (X, Y) drift was compensated during post-processing using the drift correction tool implemented in ThunderSTORM^10^, as described below. We measured the precision of the Z-drift compensation by localizing the axial position of the centroid of a dendritic segment imaged for 90 minutes. The standard deviation of the position after correction is 25 nm. The performance of the image-plane stabilization method was further evaluated during long-term DNA-PAINT acquisitions by comparing reconstructions acquired with and without axial stabilization.

### Single-molecule localization imaging

Brain slices were mounted on 5-mm-diameter round coverslips using silicone grease and positioned on the LLSM sample holder. GFP/YFP-labelled structures were targeted using the F1 docking strand and imaged with the complementary F1 imager strand conjugated to either Cy3B or ATTO 655 (Massive Photonics). GluA2 labelled with eSA-4×ALFA and an anti-ALFA single-domain antibody (NanoTag Biotechnologies, Cat. No. N1505; conjugated to the F3 docking strand) was imaged using the complementary F3 imager strand conjugated to Cy3B (Massive Photonics). Both conventional and FAST DNA-PAINT imagers were used depending on the experimental requirements. Conventional imagers were acquired with 150ms exposure times to maximize the number of photons collected per binding event and therefore localization precision, whereas FAST imagers were acquired with 50ms exposure times to shorten the overall acquisition while maintaining sufficient localization precision. DNA-PAINT acquisitions were performed using either the 560 nm or 642 nm excitation laser depending on the fluorophore coupled to the imager strand (Cy3B or ATTO655, respectively). For each fluorophore, the excitation laser was operated at full output power, corresponding to an excitation power of 4.3 and 6.2 mW measured at the back aperture of the excitation objective for the 560 nm and 642 nm excitations lasers respectively. Images were typically acquired over 128×128 or 256×256-pixel fields of view, corresponding to the region over which the light sheet provided a homogeneous illumination profile. Axial and lateral drift were compensated as described in “Image plane stabilization” section and Supplementary Materials.

### Single particle tracking imaging

For 2D-SPT, the GFP morphologic signal and single-molecule trajectories were acquired separately. For morphological imaging, a GFP z-stack comprising 100 planes with a 0.1 µm axial step size (∼10 µm axial range) was acquired over a 256×256-pixel FOV. Single-molecule imaging was subsequently performed in the same FOV for 1,000 frames with a 30 ms exposure time, using 642 nm excitation at 15–20% of the maximum laser power. For 3D SPT, the same imaging parameters were used, except that 0.1 µm RMS astigmatism at 0° was introduced with the deformable mirror through the AIO system during single-molecule acquisition to encode the axial position of individual molecules. The 642 nm excitation laser was operated at 100% of the maximum power to compensate for the reduced peak signal intensity resulting from the astigmatic PSF.

### Single-molecule localization reconstruction and quantification

For SPT imaging, single-molecule detections were localized using DAOSTORM^11^. Detection parameters were automatically optimized for each dataset using the first 15 frames of the acquisition. For 2D datasets, the expected PSF width was calculated from the microscope parameters and used to constrain PSF fitting, while the detection threshold was iteratively optimized from the fitted PSF-width distribution. For 3D datasets, axial positions were retrieved using the astigmatic PSF calibration described in “PSF engineering calibration”, within a −750 to +750 nm axial range. The detection threshold was iteratively optimized based on the stability of the fitted Gaussian parameters and axial-position distributions, assessed using a two-sample Cramér–von Mises test. For SMLM imaging, single-molecule events were localized using the ThunderSTORM plugin for Fiji^10^. Images were filtered using a B-spline wavelet filter (order 3, scale 2.0), and candidate emitters were identified as local maxima using 8-neighbourhood connectivity and an intensity threshold of 1.5 × the standard deviation of the first wavelet level. Subpixel localization was performed by weighted least-squares fitting within a 3-pixel radius, using an initial PSF sigma of 1.6 pixels. For 2D SMLM, emitters were fitted with an integrated Gaussian PSF model, whereas for 3D SMLM an elliptical Gaussian PSF model was used together with the experimentally determined astigmatic calibration described above. Multi-emitter fitting was disabled. For results visualizations, residual lateral sample drift remaining after image-plane stabilization during acquisition was corrected using the cross-correlation-based 2D drift-correction procedure implemented in ThunderSTORM. The resulting drift-corrected localizations were rendered using Normalized Gaussian rendering, with a magnification factor of 5, corresponding to a reconstruction pixel size equal to one-fifth of the calibrated sample-plane pixel size of the original acquisition. For 3D reconstructions, localizations were rendered over an axial range of −500 to +500 nm, with a 50-nm axial step.

### Localization parameters and uncertainty

For SMLM imaging, single-molecule parameters, including the number of detected localizations and lateral and axial localization uncertainties, were obtained directly from ThunderSTORM^10^ and calculated as described in the original implementation. For SPT imaging, localization uncertainties were calculated from the DAOSTORM output using a custom implementation of a previously established analytical framework for localization precision^12^, as detailed in Supplementary Materials.

### PSF engineering calibration

3D single-molecule imaging was achieved by astigmatic PSF shaping using the calibrated DM. Following AIO correction, 0.1 µm RMS of astigmatism 0° was superimposed on the corrected DM shape, generating an axially dependent ellipticity of the detection PSF. Astigmatic calibration curves were generated by axially scanning fluorescent beads through the detection focal plane over a 5 µm range with 10 nm steps. These calibration stacks were subsequently analysed using dedicated calibration procedures for SPT and SMLM imaging. For SPT imaging, bead images were fitted and linked across consecutive axial positions using the 3D-DAOSTORM z-calibration procedure. For each bead, the axial coordinate was recentered relative to its focal position, and the PSF widths along the two astigmatic axes, *W_x_* and *W_y_*, were fitted as a function of axial position according to:

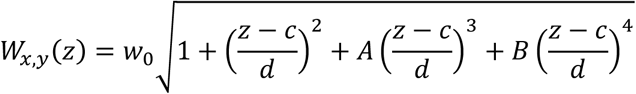

where *w*_0_, *c*, *d*, *A* and *B* were independently determined for the two astigmatic axes. The resulting calibration curves were subsequently used for 3D localization of single molecules. For each detected emitter, the PSF widths along the two axes (*σ_x_* and *σ_y_*) were determined from the fitted PSF and its axial position was estimated from the corresponding calibrated astigmatic response. For SMLM imaging, calibration stacks were analyzed using the 3D calibration procedure implemented in ThunderSTORM^10^. The relationship between the fitted PSF widths along the two astigmatic axes and the known axial bead position was used to generate a calibration curve, which was subsequently applied to estimate the axial position of individual single-molecule localizations from their measured PSF shape.

### Fourier ring correlation analysis

Fourier ring correlation analysis was performed on pairs of independently reconstructed images generated from randomly partitioned single-molecule localizations frames. First, the complete SMLM acquisition was processed in ThunderSTORM^10^, including single-molecule localization and drift correction using the full dataset. This ensured that drift estimation benefited from the complete number of detected localizations rather than being performed independently on two subsets with reduced localization density. The resulting drift-corrected localization table was exported as a CSV file and processed using a custom Python script. Localizations were first restricted to a manually defined region of interest (ROI) corresponding to the neuronal morphology. The ROI was defined in Fiji under the same spatial calibration and image dimensions as the SMLM reconstruction and saved using the Fiji ROI format. When necessary, the corresponding GFP/YFP morphological image was spatially registered to the SMLM reconstruction before ROI definition to ensure accurate correspondence between the morphological signal and the localization coordinates. Following ROI filtering, the retained localizations were randomly shuffled and divided into two non-overlapping subsets of equal size. Each subset was exported as an independent localization table and re-imported into ThunderSTORM. The two subsets were reconstructed independently using the same parameters described in “Single-molecule localization imaging”. FRC was then calculated between the two resulting super-resolution reconstructions using Fourier Ring Correlation Plugin on ImageJ and the spatial resolution was determined using the 1/7 threshold criterion. Note that the 71nm FRC resolution measured with the dendrite does not represent the fundamental limit of our method since it is also depending on the spatial frequency content of the sample itself.

### PoCA analysis

The nanoscale organization of SMLM localizations was quantified using PoCA^13^. Drift-corrected localization tables generated in ThunderSTORM were imported into PoCA for both 2D and 3D datasets. Before segmentation, localizations were restricted to the GFP/YFP-positive neuronal morphology using an ROI manually defined in Fiji by intensity thresholding of the corresponding morphological image. Localizations outside this ROI were excluded from subsequent analyses. Clusters were first identified by first-level Voronoi segmentation of the localization distribution. Objects were defined based on the local Voronoi density using an applied density factor of 4 and were retained only when containing at least 100 localizations. A second segmentation was subsequently performed within the detected clusters to identify higher-density substructures, referred to as nanodomains. This second-level Voronoi segmentation used a density factor of 0.75 and a minimum of 10 localizations per nanodomain. For each segmented object, PoCA was used to determine its dimension and the number of localizations contained within the object. For clusters, the number of nanodomains per cluster was additionally quantified. Cluster and nanodomain sizes were quantified using the principal-axis dimensions of the localization distribution, with measurements performed along two principal axes in 2D and three principal axes in 3D.

### Tracks analysis

Single-molecule localizations were linked into trajectories using TrackPy^14^ with adaptive particle linking. The linking search range was defined from a maximum molecular velocity of 5 µm.s⁻¹, corresponding to the upper range of previously reported AMPAR transport velocities^15,16^, and multiplied by a factor of 1.5 to accommodate a broad range of molecular motion while limiting spurious associations. The resulting maximum allowed displacement therefore corresponded to a velocity of 7.5 µm s⁻¹, with the frame-to-frame search radius calculated according to the acquisition time interval. Temporary disappearance of a molecule was permitted for up to 5 consecutive frames, allowing trajectories to be reconnected when detections were transiently lost. Molecular mobility was then quantified from the mean-squared displacement (MSD) of each trajectory^17,18^, from which the diffusion coefficient (D) and anomalous diffusion exponent (α), describing the magnitude and mode of molecular motion, respectively, were determined as described in Supplementary Notes.

### RESPAN segmentation

Dendrites and dendritic spines were segmented in 3D using RESPAN^19^ (Restoration Enhanced SPine And Neuron Analysis), an automated deep-learning pipeline for segmentation and quantitative analysis of neuronal morphology from fluorescence microscopy data. Before segmentation, LLSM volumes were deskewed and deconvolved using LLSpy^20^. The resulting deconvolved volumes were segmented using the pretrained nnU-Net Model 1Bv2, trained on 224 spinning-disk and Airyscan/laser-scanning confocal datasets at a reference voxel size of 65 × 65 × 150 nm and annotated to distinguish spine core and shell, dendrites, axons and soma. Input LLSM volumes were provided with their experimentally determined voxel dimensions and internally resampled by RESPAN to match the model resolution. Spine detection was restricted to objects with volumes between 0.003 and 50 µm³, while dendrite repair was enabled to reconnect fragmented dendritic segments separated by up to 2 µm. Spine-neck generation and recovery of filopodia from isolated neck fragments were enabled, together with detection of multi-headed spines. Spines with poorly supported neck assignments were filtered using a maximum neck-path length of 1 µm and a minimum nnU-Net neck-support fraction of 0.2. No image restoration was applied before segmentation.

### Spine morphological classification

Dendritic spines were classified according to their morphology using principal component analysis (PCA)^21^ followed by K-means clustering, implemented in the scikit-learn Python library^22^. PCA was performed on 26 morphological parameters extracted from the RESPAN segmentation, comprising descriptors of the entire spine, spine head and spine neck, including area, volume, surface area, length, extent, solidity, bounding-box and convex-hull measurements, as described in Supplementary Tables. The number of principal components was selected to explain 95% of the total variance, resulting in 11 principal components. K-means clustering was subsequently performed in the resulting PCA space. The number of clusters was determined using the within-cluster sum of squares (WCSS) elbow criterion and the silhouette score, both identifying seven clusters. Visual inspection identified filopodia, small mushroom, stubby, long thin, short thin, floating-head and large mushroom morphologies. Floating heads, corresponding to segmentation failures in which a spine head was detected without an associated neck, were excluded from further analysis. The remaining classes were consolidated into four morphological categories: filopodia, stubby, thin (long and short thin combined) and mushroom (small and large mushroom combined).

### Correlation of single-molecule measurements with neuronal morphology

Single-molecule measurements were spatially correlated with RESPAN-derived spine and dendrite segmentations using custom analysis based on the Open3D library^23^. For SPT datasets, point clouds representing the outer surface of the segmented neuronal morphology and the single-molecule localizations were iteratively registered by translating the localization coordinates in x, y and z. Registration was progressively refined using search ranges of ±20, ±10 and ±5 pixels, minimizing at each step the summed nearest-neighbour distance between the single-molecule localizations and the morphological surface. When no stable minimum could be identified, the original microscope-based alignment was retained. To account for the spatial uncertainty of single-molecule localization relative to the diffraction-limited morphological segmentation, individual RESPAN compartment masks were anisotropically expanded according to the median localization uncertainty of the corresponding dataset. The expansion along each dimension was determined from the localization uncertainty relative to the corresponding voxel size,

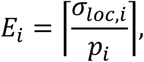

where *σ_loc_*_,*i*_ is the median localization uncertainty and *p_i_* the voxel size along dimension *i*. Expansion was performed using a spheroidal kernel, allowing the different lateral and axial localization uncertainties to be accounted for independently. Closed 3D volumes were subsequently generated from the expanded masks for each segmented spine and dendrite compartment. For SPT analysis, the occupancy of each trajectory within the expanded morphological compartments was calculated as the percentage of its localizations contained within each compartment. Tracks were assigned to a spine or dendrite when ≥95% of their localizations were contained within that compartment and <5% were contained within the alternative compartment. Tracks not fulfilling these criteria were excluded from compartment-specific analyses. For NLGN1 2D SPT, 3,836 of 14,328 trajectories were retained (26.8%), compared with 4,871 of 21,340 trajectories (22.8%) for GluA2 2D SPT and 692 of 7,019 trajectories (22.0%) for GluA2 3D SPT. For SMLM analysis, spatial assignment was performed at the level of PoCA-defined clusters and nanodomains rather than individual localizations to reduce computational load. The centroid of each cluster or nanodomain was calculated from its constituent localizations, and structures were assigned to individual spines when their centroid was contained within the corresponding anisotropically expanded RESPAN volume. Of the 853 clusters initially identified by PoCA, 476 were assigned to segmented morphological compartments. After excluding ambiguous assignments to multiple spines, 432 clusters were retained for subsequent analysis, corresponding to 50.6% of the initial population.

### Statistical analysis

All reported results are derived from at least three independent experiments to ensure replicability. No statistical test was used to predetermine sample sizes. Statistical analysis and data plotting were performed with Prism 11.1.0 (GraphPad). Datasets were analyzed with the Shapiro-Wilk test for normality, and parametric (P > 0.05) or nonparametric (P < 0.05) statistical tests were performed as appropriate. Data was analyzed with the robust regression and outlier removal (ROUT) method to identify outliers. Data is presented as Mean ± SEM or Median ± standard deviation unless otherwise stated. Test details and statistical outcomes for all experiments are reported in the relevant figure legends.

## Code avaibility

Data analysis for SPT, including single-molecule detection, localization, trajectory reconstruction, and PCA-based classification, was performed using custom Python scripts combining existing open-source packages. These scripts are available from the corresponding authors upon request. SMLM analysis was performed using the open-source software ThunderSTORM^10^ and PoCA^13^, while dendritic spine segmentation was performed using the open-source software RESPAN^19^.

## Data availability

Other data that support the findings of this study are available from the corresponding authors upon request.

## Acknowledgements

We thank P. Costet, C. Martin at the "Breeding core-facility Exempte d’Organisme Pathogène Spécifique" (EOPS) of the common animal facility services of Bordeaux for breading of the mouse strain. We thank H. El Oussini at the "Core-Facility In Vivo experimental" (PIV-EXPE) of the Bordeaux Neurocampus for maintenance of the mouse strain and completion of the experimental and surgical tests. We thank A. Caralp and people involved within the Cell Biology Facility from Bordeaux Neurocampus for their assistance with organotypic brain slice preparation and culture, C. Breillat for her contribution to the development of the AP-GluA2 mouse line. The microscopy was done in the Bordeaux Imaging Center a service unit of the CNRS-INSERM and Bordeaux University, member of the national infrastructure France BioImaging supported by the French National Research Agency (ANR-10-INBS-04). We thank the Howard Hughes Medical Institute, Janelia Research Campus for providing all the technical information to reproduce the lattice light-sheet microscope upon signature of a research license agreement. We thank J. Legrand and C. Veilly from Imagine Optic for their contributions to the development of the Active Image Optimization (AIO) software and its optical implementation. We also thank F. Levet for his assistance with PoCA for protein nano-organization quantification, and L. Hammond for his assistance with RESPAN for dendritic spine segmentation. This work was supported by the Conseil Régional de Nouvelle-Aquitaine (AAPPF2020I-2019-8336710), the Bordeaux Neurocampus core facilities (ANR-10-LABX-43), by European Research Council (ERC) grants to D.C. (339541 and 787340), received financial support from the French government in the framework of the University of Bordeaux’s IdEx "Investments for the Future" program / GPR BRAIN_2030 (ANR-23-CE16-0019-03 and ANR-23-CE11-0007-02) to M.S., from Institut National de la Santé et de la Recherche Médicale (Crédit Equipement 2020 EQT-313831) to M. D. A.M.G. was supported by the Canadian Institutes of Health Research (CIHR; 158090) and the University of Bordeaux Initiative of Excellence (IdEx) postdoctoral fellowships. M.M. was supported by a CIFRE doctoral fellowship funded by the Association Nationale de la Recherche et de la Technologie (ANRT; 2020/0147), a postdoctoral fellowship from the Grand Programme de Recherche (GPR) BRAIN_2030 of Bordeaux Neurocampus, and postdoctoral funding from Vrije Universiteit Amsterdam.

## Author contributions

M.Mal., M.Mon., M.S., D.Ch., M.D., A.M.G. conceived the idea and designed the experiments. M.S., D.Ch., M.D., A.M.G. supervised the research. M.Mal., M.D., B.D., M.Mar., F.H. developed AIO-SMLM-LLSM imaging. M.S., C.L., P.L.S-J., developed and produced mSA and eSA labelling probes. O.T. developed and provided the AP-Nlg1 mouse line. M.Mal., C.C.G., M.Mon., A.M.G. prepared samples and performed experiments. B.A.A.J.W.W., M.Mal., C.C.G., M.Mon., D.Cri., B.D. analyzed the data. M.Mal. wrote the paper with input from all authors. All authors reviewed and approved this version of the manuscript.

## Supplementary Material

### Supplementary Notes

#### Image plane stabilization experiment

To illustrate the clear benefit of our method, we acquired 40,000 DNA-PAINT frames of a GFP+ dendrite with and without axial stabilization (Fig. S2b-c). The full acquisition lasted 90 minutes, during which a 10 µm-thick GFP-channel stack was acquired every 2:15 minutes to measure and correct spatial drift via cross-correlations. The DNA-PAINT acquisition plane stabilized on a dendrite at the middle plane of the first GFP stack (Fig. S2a, bottom row). DNA-PAINT reconstruction (Fig. S2c, bottom left) was further corrected for lateral drift using ThunderSTORM’s drift-correction algorithm (Fig. S2c, bottom right). Without axial drift compensation, the imaged plane is not stabilized on the dendrite (Fig S2b, to row), and the raw DNA-PAINT reconstruction consists of localizations spanning several axial positions (Fig. S2c, top left). As expected, post-processing lateral-drift correction cannot recover an accurate dendritic reconstruction (Fig. S2c, top right).

#### DAOSTORM detection parameters

For SPT analysis, DAOSTORM detection parameters were automatically optimized independently for each dataset using the first 15 frames of the acquisition. Starting from a low detection threshold, the threshold was iteratively increased until predefined criteria based on the properties of the detected PSFs were satisfied. Different optimization criteria were used for 2D and astigmatic 3D acquisitions.

For 2D SPT, the theoretical diffraction-limited PSF size was first estimated from the microscope parameters. The theoretical FWHM was calculated as:

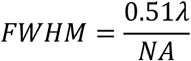

where *λ* is the emission wavelength and NA is the numerical aperture of the detection objective. The corresponding Gaussian width was calculated as:

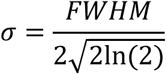

and converted into camera-pixel units according to:

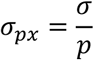

where *p*is the calibrated sample-plane pixel size. The resulting theoretical PSF width was used to define the DAOSTORM “sigma_range” as:

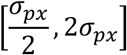

At each iteration, the distribution of fitted PSF widths was evaluated. Detections with fitted widths below the lower outlier boundary:

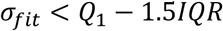

were considered inconsistent with the expected single-molecule PSF, where *Q*_1_is the first quartile and IQR = *Q*_3_ − *Q*_1_. The detection threshold was progressively increased until no such low-width outliers remained, and the resulting threshold was subsequently applied to the complete dataset.

For 3D SPT, the PSF widths along the two astigmatic axes (*σ_x_* and *σ_y_*) vary with axial position and therefore could not be constrained using the 2D criterion. DAOSTORM was initialized with an axial localization range of −750 to +750 nm relative to the focal plane, corresponding to approximately 1.5 times the light-sheet thickness. At each detection-threshold iteration, the distributions of *σ_x_*, *σ_y_*and reconstructed *z*positions were compared with those obtained at the preceding threshold. A two-sample Cramér–von Mises test was independently applied to each distribution to assess whether the fitted localization parameters changed significantly. The detection threshold was retained once the distributions of *σ_x_*, *σ_y_*and *z*no longer changed significantly, indicating that the detection parameters had reached a plateau.

#### Localization uncertainty estimation for SPT

Localization uncertainties for DAOSTORM detections were calculated using a custom implementation based on the analytical maximum-likelihood estimator (MLE) approximation described by (Rieger & Stallinga, 2014) incorporating the finite pixel size and the contribution of background noise.

For 2D localizations, the noise parameter *τ*_2*D*_was calculated as:

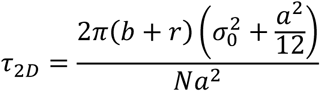

and the lateral localization uncertainty was estimated as:

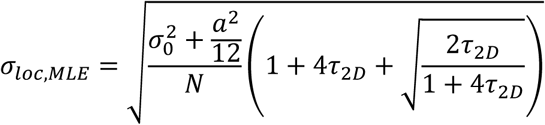

where *σ*_0_ is the fitted PSF width, *a* is the effective camera pixel size, *N* is the number of detected photons, and *b* and *r* account for the background and camera-noise contributions, respectively.

For 3D astigmatic localizations, the same framework was extended to account for the different fitted PSF widths along the two lateral axes, *σ*_0_*_x_* and *σ*_0_*_y_*. The corresponding noise parameter was calculated as:

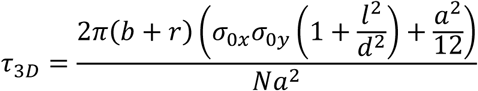

where *σ*_0*x*_ and *σ*_0*y*_ represent the standard deviation of the fitted PSF, 2 *l* the distance between focal planes *d* the measure of astigmatic focal depth, *a* the in-plane pixel size, *N* the integrated signal photon count, *b* the integrated background photon count, and *r* the read-noise variance. We used g=1, since recordings were made using a sCMOS detector and *r* = 0, as localization uncertainties were estimated from fitted total signal and background photon counts for each localization and detector read noise was not modelled separately. The resulting parameters were used to estimate the lateral and axial localization uncertainties for each detected molecule according to the (Rieger & Stallinga, 2014) framework.

The lateral localization uncertainties were calculated as:

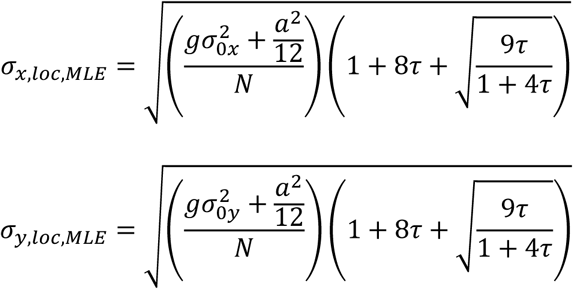

where *σ*_0*x*_ and *σ*_0*y*_ are the standard deviation of the fitted PSF, *a* is the in-plane pixel size, *N* is the integrated signal photon count, *g* isthe excess noise factor, and *τ* isthe scaled background parameter defined above.

The axial localization uncertainty was subsequently obtained by propagating the lateral localization uncertainties through the astigmatic PSF calibration. The calibration-dependent quantity *F* was defined as:

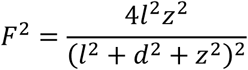

and its uncertainty as:

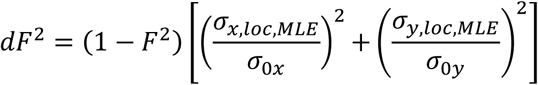

Finally, the axial localization uncertainty was calculated as:

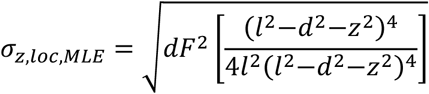

where *z* is the fitted axial centre position of the PSF, 2 *l* is the distance between focal planes, and *d* is the measure of astigmatic focal depth. Thus, localization uncertainties were determined independently for the two lateral dimensions and subsequently propagated through the astigmatic calibration to estimate the axial localization uncertainty for each 3D localization.

#### Single-molecule dynamics quantification

Molecular mobility was quantified from the mean-squared displacement (MSD) of each trajectory according to (Kepten et al., 2015; Michalet, 2010) :

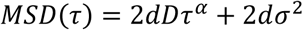

where *D* is the diffusion coefficient, *τ* the time lag, *α* the anomalous diffusion exponent, *σ* the localization uncertainty and *d* the number of spatial degrees of freedom. As the analysed receptors were confined to the neuronal plasma membrane, *d* = 2 was used for both 2D and 3D localization datasets. The diffusion coefficient *D* was estimated by linear fitting of the first 4 MSD lag points, while accounting for the intercept associated with localization uncertainty:

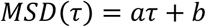

with,

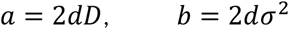

The diffusion coefficient was therefore calculated as:

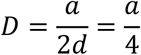

The anomalous diffusion exponent *α*was determined by fitting the MSD in logarithmic space. Neglecting the localization-uncertainty contribution after correction, the anomalous diffusion model can be expressed as:

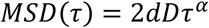

Taking the logarithm gives:

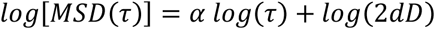

Thus, *α* corresponds to the slope of the linear regression of log[MSD(*τ*)] against log(*τ*):

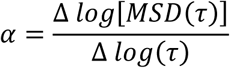

The fit was performed using a maximum of the first 10 MSD lag points; for shorter trajectories, the number of fitted points was limited to 50% of the available MSD points.

### Supplementary Figures

**Fig. S1.**
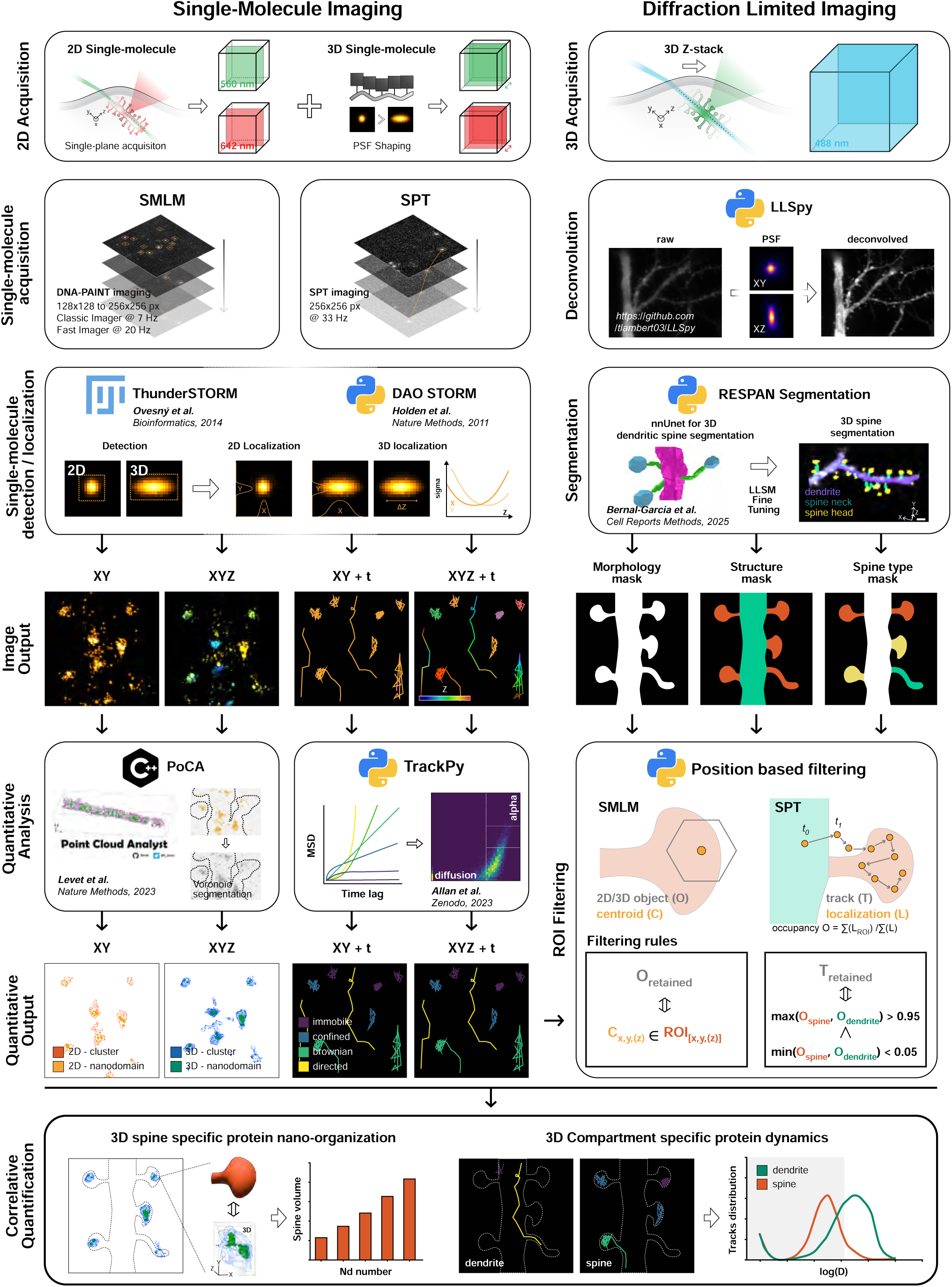
ALPINE: an analysis pipeline for cellular contextualization of endogenous nanoscale proteins in tissue. Integrated imaging and analysis pipeline combining single-molecule (left) and diffraction-limited (right) imaging acquired from the same FOV. Single-molecule acquisitions are used to quantify the nanoscale organization of endogenous proteins by DNA-PAINT and protein dynamics by SPT. In parallel, diffraction-limited 3D images are deconvolved and used to segment dendritic spines through a deep-learning-based approach, providing morphological and structural 3D masks. Reconstructed molecular clusters and trajectories are then spatially filtered according to their position within the segmented cellular masks. This integrated analysis enables compartment-specific quantification of protein nano-organization and dynamics and contextualizes single-molecule information with respect to spine morphology and cellular architecture.

**Fig. S2.**
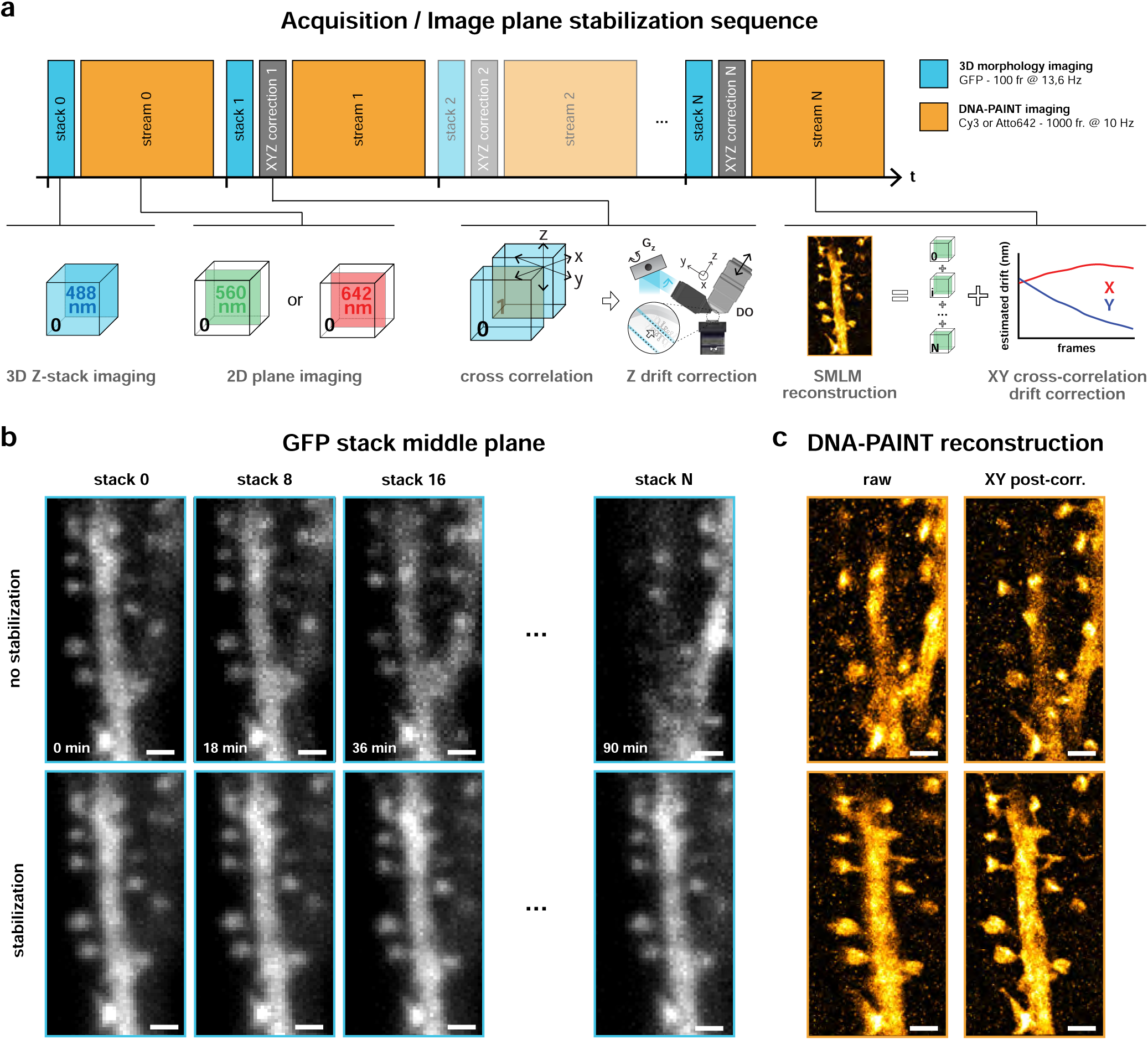
Interleaved image-plane stabilization enables long-term SMLM acquisition without XYZ drift. **a,** Image acquisition sequence for stabilized SMLM imaging. SMLM acquisition is divided into 2D sub-acquisitions (stream), interleaved with 3D stack imaging of the morphological signal (stack). After each acquisition interval, the newly acquired 3D GFP stack is compared with the previous stack by cross-correlation to determine the Z shift. The calculated Z offset is applied to the galvanometer Z (Gz) and detection objective (DO), respectively, to move light-sheet and detection plane and compensate for axial drift. The individual SMLM sub-acquisitions are then combined to generate the full SMLM acquisition and reconstruction. Finally, XY cross-correlation is applied to compute and correct lateral drift. **b,** Representative middle planes of GFP stacks illustrating the effect of image-plane stabilization over a 90-min acquisition. Without stabilization (top), progressive XYZ drift is observed between consecutive GFP stacks. Image-plane stabilization (bottom) maintains the imaged region and focal plane throughout the acquisition while a small drift persists. **c,** SMLM reconstructions of the same FOV as in **b**, acquired without (top) or with (bottom) image-plane stabilization. Image-plane stabilization corrects XYZ drift during acquisition. Subsequent XY post-correction by cross-correlation of the SMLM reconstruction compensates for lateral drift, improving reconstruction accuracy. Note the similarity between the XYZ drift-corrected DNA-PAINT reconstruction (**c,** bottom right) and the initial GFP stack middle plane (**b**, bottom left), showing that the initial ROI is maintained throughout the acquisition. In contrast, without image-plane stabilization, the DNA-PAINT reconstruction (**c**, top right) combines SMLM localizations acquired at different planes due to XYZ drift and therefore does not faithfully represent the initial ROI (**b**, top left). **Scale bar,** 1 μm (**b, c**).

**Fig. S3.**
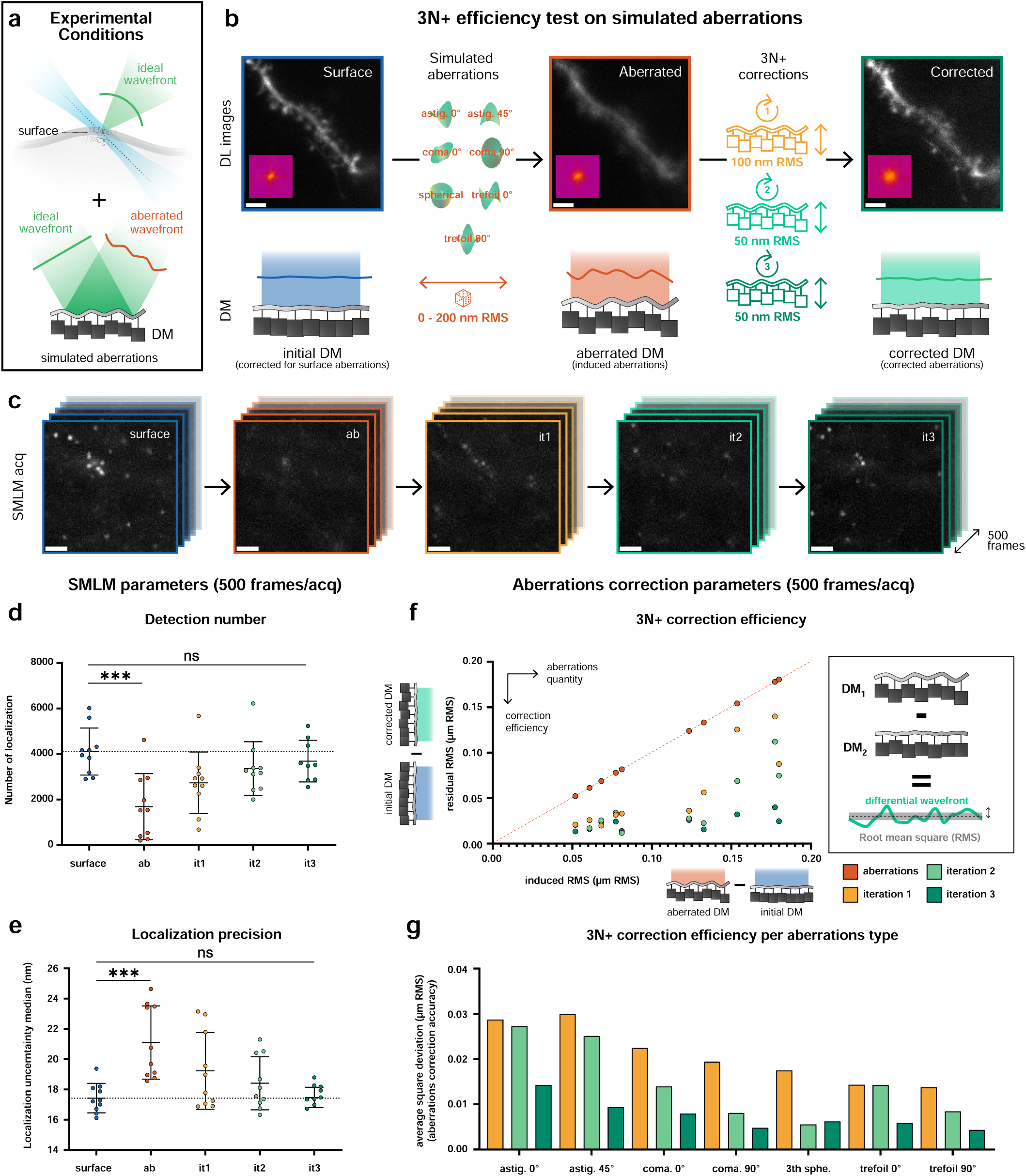
3N+ optical aberrations correction efficiency on simulated aberrations in tissue. **a,** Schematic representation of the experimental conditions. Imaging is performed at the surface of the brain tissue, where optical aberrations are minimal. Simulated aberrations, described by individual Zernike modes, are introduced using the deformable mirror (DM). **b,** Evaluation of 3N+ correction efficiency on simulated aberrations. Blue, a FOV containing a representative morphological structure is selected and the DM is initially calibrated to provide an aberration-free reference. Red, image quality is degraded by applying individual Zernike aberrations with amplitudes ranging from 0 to 200 nm RMS to the DM. Green, the induced aberrations are corrected over three consecutive 3N+ iterations using decreasing correction amplitudes (100, 50 and 50 nm RMS), progressively restoring image quality. **c,** Effect of induced optical aberrations and their correction on SMLM acquisition in the same FOV as in **b**. Representative SMLM acquisitions are shown at the surface, after application of the simulated aberration (red), and after each of the three consecutive 3N+ correction iterations (yellow green). **d,** Number of single-molecule detections measured at each experimental step over 500-frame acquisitions. Simulated aberrations reduce the number of detected molecules, which progressively recovers over the three consecutive 3N+ correction iterations. **e,** Median localization uncertainty measured at each experimental step over 500-frame acquisitions, showing recovery toward surface-level performance following 3N+ correction. **f,** Effect of 3N+ correction for the acquisitions shown in **d** and **e**. Inset, induced and residual aberration RMS values are determined from the DM actuator positions and calibration matrix by computing the corresponding differential wavefront. Residual aberrations are plotted against the initially induced aberrations, showing efficient correction across a range of aberration amplitudes after three 3N+ iterations. **g,** Aberration-specific analysis of 3N+ correction for the acquisitions shown in **d-f**. The DM actuator positions and calibration matrix are used to retrieve the correction amplitude associated with each Zernike aberration mode, enabling evaluation of the correction efficiency for individual aberration types.

**Table S1.** Morphological parameters used for PCA-based spine classification. Twenty-six morphological parameters extracted from RESPAN segmentations were used for principal component analysis (PCA). Parameters describe the complete spine and its head and neck compartments.

| Compartment | RESPAN parameter | Description | Unit |
| --- | --- | --- | --- |
| Whole spine | spine_area | Cross-sectional or projected area of the entire spine | $\mu\text{m}^2$ |
| Whole spine | spine_vol | Volume of the entire spine | $\mu\text{m}^3$ |
| Whole spine | spine_surf_area | Surface area of the entire spine | $\mu\text{m}^2$ |
| Whole spine | spine_length | Total morphological length from the dendrite base to the spine-head tip | $\mu\text{m}$ |
| Whole spine | spine_length_euclidean | Straight-line (Euclidean) distance from the distal spine tip to its attachment point at the dendrite | $\mu\text{m}$ |
| Whole spine | spine_bbox_vol | Volume of the tightest image-axis-aligned 3D bounding box enclosing the spine mask | $\mu\text{m}^3$ |
| Whole spine | spine_extent | Ratio of spine volume to bounding-box volume (spine_vol / spine_bbox_vol) | Dimensionless |
| Whole spine | spine_solidity | Ratio of spine volume to convex-hull volume (spine_vol / spine_convex_vol) | Dimensionless |
| Whole spine | spine_convex_vol | Volume of the convex hull enclosing the entire spine | $\mu\text{m}^3$ |
| Spine head | head_area | Cross-sectional or projected area of the segmented spine head | $\mu\text{m}^2$ |
| Spine head | head_vol | Volume of the spine head | $\mu\text{m}^3$ |
| Spine head | head_surf_area | Surface area of the spine head | $\mu\text{m}^2$ |
| Spine head | head_length | Morphological length of the spine head from the neck-head boundary to the head tip | $\mu\text{m}$ |
| Spine head | head_bbox_vol | Volume of the tightest image-axis-aligned 3D bounding box enclosing the spine-head mask | $\mu\text{m}^3$ |
| Spine head | head_extent | Ratio of spine-head volume to its bounding-box volume (head_vol / head_bbox_vol) | Dimensionless |
| Spine head | head_solidity | Ratio of spine-head volume to its convex-hull volume (head_vol / head_convex_vol) | Dimensionless |
| Spine head | head_convex_vol | Volume of the convex hull enclosing the spine head | $\mu\text{m}^3$ |
| Spine head | head_convex_hull_ratio | Ratio describing how tightly the spine head conforms to its convex hull relative to other structural boundaries | Dimensionless |
| Spine neck | neck_area | Cross-sectional or projected area of the segmented spine neck | $\mu\text{m}^2$ |
| Spine neck | neck_vol | Volume of the spine neck | $\mu\text{m}^3$ |
| Spine neck | neck_surf_area | Surface area of the spine neck | $\mu\text{m}^2$ |
| Spine neck | neck_length | Morphological length of the spine neck from its origin to the neck-head boundary | $\mu\text{m}$ |
| Spine neck | neck_bbox_vol | Volume of the tightest image-axis-aligned 3D bounding box enclosing the spine-neck mask | $\mu\text{m}^3$ |
| Spine neck | neck_extent | Ratio of spine-neck volume to its bounding-box volume (neck_vol / neck_bbox_vol) | Dimensionless |
| Spine neck | neck_solidity | Ratio of spine-neck volume to its convex-hull volume (neck_vol / neck_convex_vol) | Dimensionless |
| Spine neck | neck_convex_vol | Volume of the convex hull enclosing the spine neck | $\mu\text{m}^3$ |

